# Molecular, anatomical, and functional organization of lung interoceptors

**DOI:** 10.1101/2021.11.10.468116

**Authors:** Yin Liu, Lucas Kinsey, Alex J. Diaz de Arce, Mark A. Krasnow

## Abstract

Interoceptors, sensory neurons that monitor internal organs and physiological states, are essential for regulating physiology, shaping behavior, and generating internal perceptions. Here, we present a comprehensive transcriptomic atlas of mouse lung interoceptors, identifying 10 molecular subtypes. These subtypes differ in developmental origin, sensory receptor repertoire, signaling molecules, anatomical receptive fields, terminal morphologies, and cell contacts. Activity recordings and functional interrogation of two *Piezo2*^+^ subtypes revealed distinct sensory properties and separate roles in breathing control: one regulates inspiratory time; the other regulates inspiratory flow. Together, these findings suggest that this pronounced cellular diversity of lung interoceptors enables the system to encode diverse and dynamic sensory information, mediate myriad local cellular interactions, and regulate respiratory physiology with precision.

## INTRODUCTION

As we vividly perceive the external world through the five canonical senses, the nervous system also continuously monitors our internal organs and environment ^1–4^. This internal monitoring is mediated by neurons that detect signals associated with internal states, dubbed "interoceptors" over a century ago ^5^. In mammals, a major fraction of interoceptors directly innervate internal organs and are pseudounipolar cells with their cell bodies located in cranial or dorsal root ganglia. For each organ interoceptor, one axonal branch projects to the target organ(s), where its terminals detect various signals, while the other projects to the brainstem or spinal cord, relaying sensory information to the central nervous system via synapses with second-order neurons. The sensory information conveyed by these neurons is essential for maintaining physiological homeostasis in processes such as digestion, circulation, and breathing ^6–8^, for triggering protective reflexes such as vomiting and coughing ^9,10^, and for generating interoception, the higher-order brain representation of internal states ^4,11^.

The mammalian lungs are richly innervated by interoceptors whose cell bodies, at least in mice, reside predominantly in the sensory ganglion of the tenth cranial nerve, the vagus nerve (Figure S1B). The vagal ganglion contains neurons that innervate most internal organs, with approximately 10-20% projecting to the lungs (these neurons were commonly referred to as bronchopulmonary sensory neurons but abbreviated pulmonary sensory neurons, or PSNs, here). Classical electrophysiological studies on PSNs have identified three major classes: two classes of myelinated, fast-conducting A-fiber neurons that both responding to lung inflation but differ in adaption properties (rapidly adapting receptors, RARs; slowly adapting receptors, SARs), and one class of unmyelinated, slow-conducting C-fibers that respond to harmful exogenous and endogenous chemicals such as irritants and inflammatory factors ^12–14^. Additional diversity likely exists, as PSNs with other physiological properties (e.g., Aδ high threshold receptors (AHTRs) and deflation receptors^15,16^) and distinct developmental origins (e.g., C-fibers derived from neural crest vs neural placode ^17^) have been described. PSN terminals range from free-nerve endings to specialized structures associated with smooth muscle-associated and neuroepithelial bodies (NEBs) ^18^. Mechanical and pharmacological challenges to the lungs, which activate different PSN populations, can evoke diverse respiratory responses, including changes in breathing patterns, bronchial tone, and airway secretion ^8,14,19^. Although each modality has been extensively characterized, integrating these observations within the same neuronal populations has been challenging, owing to the incompatibility of experimental approaches and the lack of subtype-specific manipulation tools.

Over the past decade, advances in molecular profiling have revealed substantial heterogeneity among vagal sensory neurons, and genetic strategies for targeting specific subpopulations of these neurons have proven powerful for linking results from different experimental modalities ^20–25^. Several studies have combined retrograde labeling from the internal organs with transcriptomic profiling to delineate the organization of interoceptors innervating specific organs, including the lungs ^25–27^. Yet, it remains unclear whether the full molecular diversity of PSNs has been uncovered. Furthermore, no single molecularly defined PSN subtype has been comprehensively characterized across multiple dimensions integrating its molecular signature, anatomical distribution, sensory properties, and physiological functions.

Here, we present a comprehensive transcriptomic atlas of mouse PSNs, defining 10 molecular subtypes and elucidating their relationships to classical physiological classes. We identify robust molecular markers for each subtype, delineate their sensory repertoire, and characterize their distinct neurotransmitter and neuropeptide usage. Genetic labeling of selected subtypes reveals striking differences in terminal distribution, morphology, and cellular contacts within the lungs. Functional interrogation of two *Piezo2*^+^ subtypes demonstrates their divergent sensory properties and exquisitely specific homeostatic roles in breathing regulation. Together, these findings reveal the remarkable complexity of sensory neurons within a single organ and the precise contributions of individual molecular subtypes to organ-specific physiological control.

## RESULTS

### Labeling and single-cell mRNA profiling of vagal PSNs

To generate a comprehensive gene expression atlas of PSNs, we performed single-cell RNA sequencing (scRNA-seq) on vagal sensory neurons that innervate the mouse lung (Figure 1A). Vagal PSNs were labeled by intratracheally instilling wheat germ agglutinin (WGA) with fluorescent conjugates into the lungs and allowing 3-7 days for WGA uptake and retrograde transport to the cell bodies in the ganglion. This approach labeled 431±48 neurons per ganglion (mean±SD, n=6 ganglia), ∼15% of the ganglion’s ∼3000 neurons ^28^. In contrast, only a smaller number of dorsal root ganglion neurons were labeled (Figure S1A), which we did not focus on in this study. Co-instillation of a second WGA conjugated to a different fluorophore labeled an almost identical population of vagal ganglion neurons (95% overlap), whereas co-injection of the second WGA into the stomach labeled a spatially intermingled but entirely non-overlapping (0% overlap) population (Figure 1B and C), indicating that PSN labeling by this method is efficient and specific. We also compared labeling efficiency between WGA and adeno-associated virus (AAV) instillation, which was previously used to sequence vagal sensory neurons innervating multiple internal organs, including the lungs. WGA labeling yielded proximately twice as many labeled neurons as AAV (Figure 1D), suggesting that this method is likely to target a more comprehensive set of vagal PSNs.

**Figure 1.**
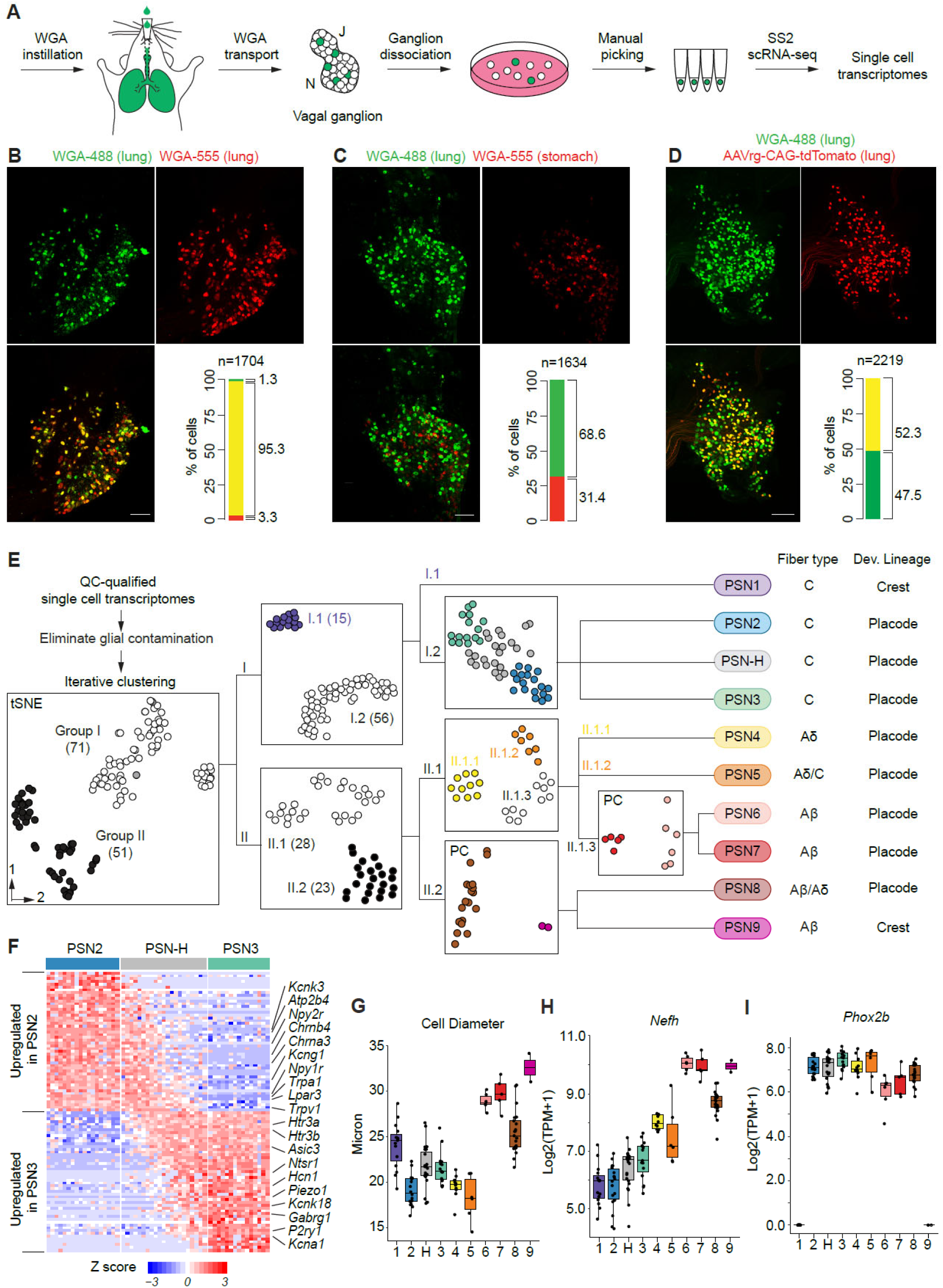
Labeling, transcriptomic profiling, and identification of 10 molecular subtypes of vagal PSNs. (A) Strategy for scRNA-seq of vagal PSNs. PSNs were labeled by intratracheal instillation of a fluorescent WGA conjugate (green). Vagal ganglia (J, jugular ganglion; N, nodose ganglion) were dissociated, and WGA-labeled neurons were individually picked and placed in separate tubes. scRNA-seq was carried out in each tube using the Smart-seq2 (SS2) method. (B, C) Retrograde labeling with two fluorescent WGA conjugates (WGA-488, green; WGA-555, red) as indicated for vagal sensory neurons from the lungs (n=3 mice) or from the lungs and the stomach (n=2 mice). Images are maximum projections of the entire ganglia after BABB clearance, with quantifications at bottom right. n, total labeled neurons scored (both fluorescent labels). (D) Retrograde labeling with WGA-488 and AAVrg-CAG-tdTomato instilled into the lungs (n= 3 mice). AAV was instilled three weeks prior to WGA-488 instillation. (E) Computational cell clustering pipeline for subtype identification based on similarity of scRNA-seq expression profiles. In each round of iterative clustering (hierarchical steps, left to right), cells were clustered into two groups based on highly variable genes, and only neurons that stably clustered together were promoted to the next round (each intermediate cluster numbered as indicated, e.g., I.1). Iteration ended when no statistically significant separation was generated, or the number of cells in one group was <5. Intermediate clusters are shown in black or white (gray cells did not stably cluster), and cells in the final clusters are color coded and their subtype designation given at right along with predicted fiber type and developmental origin from results in panels G-I. The number of neurons in each cluster is indicated in parentheses. All plots are tSNE projections except two marked with PC, which show only the first two principal components of PCA because no more than two PCs were statistically significant. (F) Heatmap showing the expression pattern of genes that are differentially expressed between PSN2 and PSN3 neurons across PSN2, H and 3. Many ion channels and GPCRs (indicated) exhibit a gradient of expression in PSN-H neurons, suggesting they are functional intermediates. (G-I) Cell diameters (G) and expression levels of *Nefh* (H, correlates with myelination states) and *Phox2b* (I, developmental lineage marker) of neurons of each PSN subtype. Dots represent values for individual neurons, and hinges correspond to 25^th^ and 75^th^ percentiles.

Single-cell suspensions from labeled vagal ganglia were visualized under a fluorescence microscope, where individual labeled PSNs were imaged, manually picked, and transferred into individual tubes for cDNA synthesis. This approach minimized cell damage and avoided size-based biases introduced by fluorescence-activated cell sorting (FACS). Neural crest-derived PSNs constitute only ∼7% of the total population (Figure S1B and C) and exhibited a lower survival rate following dissociation compared to neural placode-derived PSNs. To ensure adequate representation of neural crest-lineage neurons in our dataset, we performed WGA labeling in animals that allowed us to distinguish neural crest-vs. neural placode-derived neurons based on tdTomato reporter expression during cell selection (*Wnt1-Cre;Ai14/+*, neural-crest specific; *CGRPα^CreERT^*^2^*^/+^;Ai14/+*, neural crest-enriched; *Phox2b-Cre;Ai14/+*, neural placode specific, Figure S1B and C) ^29–31^. cDNA synthesis and RNA amplification from individual neurons were performed using the SmartSeq2 protocol, with fewer amplification cycles (14–15) than the standard (18–21) to minimize mRNA amplification bias. We successfully obtained sufficient cDNA for sequencing library construction from 136 labeled cells, representing approximately one-third of the total PSNs in a ganglion.

Sequencing reads were aligned to the mouse transcriptome (RefSeq), and cells with fewer than 2 million mapped paired-end reads or fewer than 10,000 expressed endogenous genes were excluded. To remove glial-contaminated cells, we performed principal component analysis (PCA) using genes enriched in satellite glial cells from peripheral sensory ganglia ^32^ (Figure S1D and E). The final dataset comprised 123 high-quality neuronal expression profiles, with an average of 5.4 million total paired-end reads per cell (range: 2.4–12.4 million), mapping to an average of 12,500 endogenous genes (range: 10,300–14,000 genes). The number of expressed genes detected per cell was 60–440% higher than in previous droplet-based scRNA-seq datasets of vagal sensory neurons (average: 2,300–8,000 genes per cell) ^21–23,25,27^. To assess dataset quality, we examined the expression levels of housekeeping genes (*Gapdh* and *Actb*), the vagal sensory neuron marker *Slc17a6*, and the fluorescent marker gene *tdTomato*. All were consistently detected across cells, with no dropout and minimal variance (Figure S1F and G). These results indicate that our vagal PSN scRNA-seq dataset is of high quality, providing a robust resource for transcriptomic analysis.

### Identification of 10 molecular subtypes of vagal PSNs

To identify molecularly distinct subtypes, we clustered PSNs based on the similarity of their expression profiles using a customized iterative strategy adapted from one developed for mouse cortical neurons ^33^ (Figure 1E, Materials and Methods). This approach revealed eight clusters, designated as PSN subtypes (PSN1 through PSN8). We also identified two additional PSN subtypes: PSN9 and PSN-H. PSN9, represented by only two neurons, was clearly separated from PSN8 during clustering and was validated via in situ hybridization as a rare but distinct subtype (Figure 1E and 2D). PSN-H neurons appeared as molecular hybrids of PSN2 and PSN3, co-expressing many of the differentially expressed genes that distinguish these subtypes. Depending on the subset of highly variable genes used for clustering, PSN-H cells variably grouped with either PSN2 or PSN3 (Figure 1E and F). We did not identify any uniquely expressed genes that distinguish PSN-H from PSN2 and PSN3. Instead, most co-expressed genes, including many receptors and channels, exhibited a gradual transition in expression levels across the population (Figure 1F), suggesting that PSN-H represents an intermediate functional state between PSN2 and PSN3.

**Figure 2.**
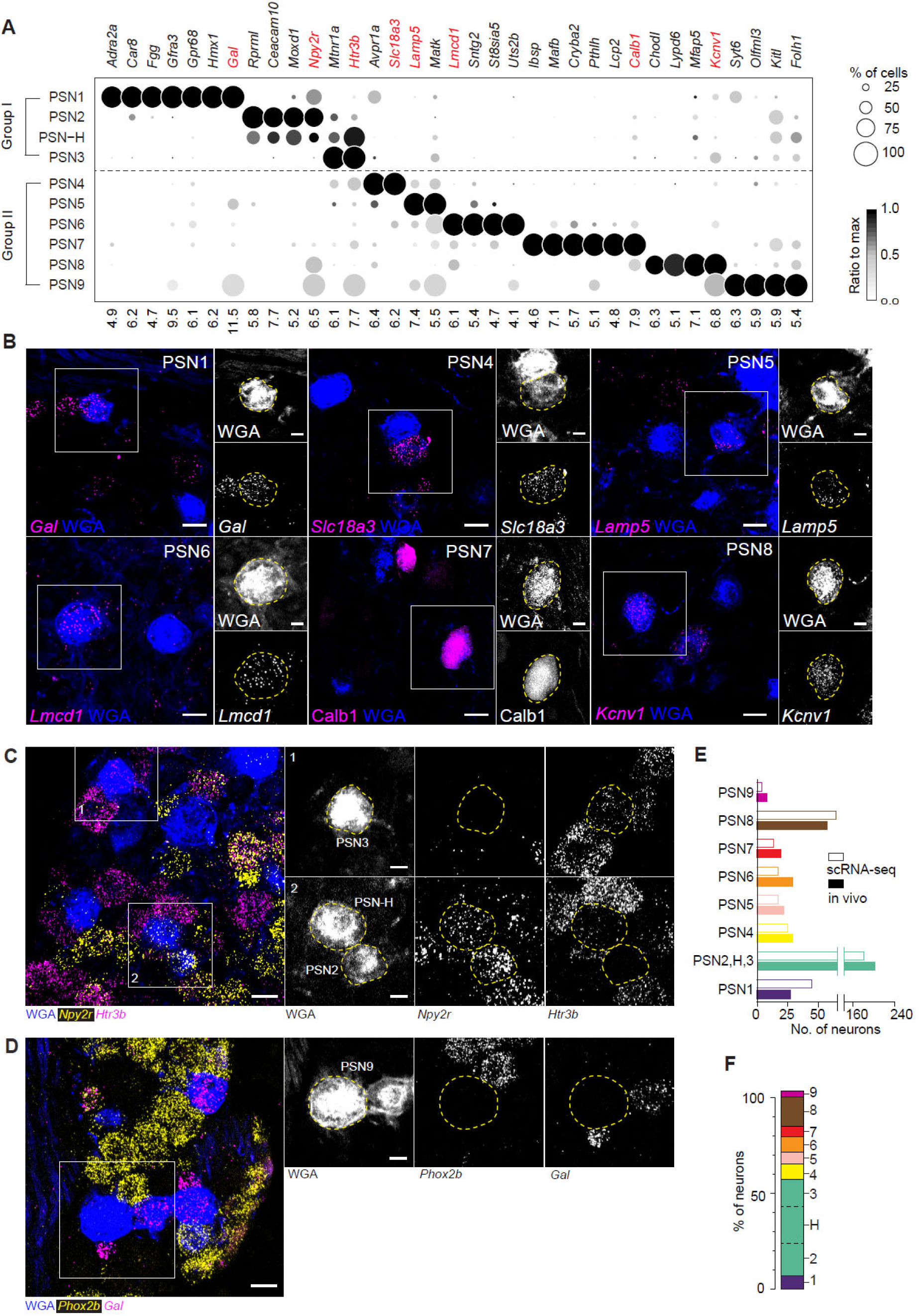
Molecular markers and abundances of PSN subtypes. (A) Dot plot showing expression levels of PSN subtype marker genes. Numbers at bottom are mean log_2_(TPM+1) values of the subtypes with the highest average expression levels of the indicated genes, with a threshold set at TPM>=1 (TPM, transcript per million). This format is used for expression dot plots in all figures, unless noted otherwise. Genes in red were used for validation analysis by in situ hybridization (ISH) and immunostaining (B-D). Group I and II correspond to the two clusters generated in the first round of iterative clustering shown in Figure 1E. (B-D) ISH or immunostaining of the indicated genes on mouse vagal ganglion sections with WGA retrograde labeling from the lungs. Close-ups of boxed areas are shown in insets with channels split. Scale bars: 20 µm and 10 µm (insets). (E) Estimated number of PSN neurons of each subtype in one vagal ganglion based on ISH/immunostaining results (solid bars) or sequencing results (open bars). Values were scaled to a total of 400 PSNs per ganglion. PSN2, H and 3 are plotted together because no unique marker is identified for PSN-H neurons. Quantification sample sizes are given in Figure S2D. (F) Estimated proportions of individual PSN subtypes. All subtypes except PSN2, H, and 3 are estimated based on ISH/immunostaining results. PSN2, 3 and H are estimated based on sequencing results (divided by dashed lines). Sum of the estimated proportions totals 105%, indicating the 10 subtypes account for all PSNs labeled by our strategy.

The traditional classification of PSNs is based in part on their fiber types, which correlate with cell body diameter and neurofilament heavy chain expression levels ^34^. PSN1, 2, 3, and H – grouped together as "Group I neurons" in the initial round of clustering — consist of small to medium diameter neurons that express low levels of *Nefh* (Figure 1G and H), suggesting they are unmyelinated C-fiber neurons. PSN4 through 9 (Group II neurons) include small (PSN4 and 5), medium (PSN8), and large diameter neurons (PSN6, 7 and 9) (Figure 1G). Among these, PSN6, 7 and 9 express high levels of *Nefh*, suggesting they are thickly myelinated (Aβ-fiber) neurons. PSN8 neurons express intermediate levels of *Nefh*, suggesting they are myelinated but likely with thinner myelination (possibly Aβ or Aδ-fiber). PSN4 and 5 express slightly higher levels of *Nefh* than Group I neurons, suggesting they may be thinly myelinated (Aδ-fiber) or unmyelinated (C-fiber) (Figure 1H). We also inferred developmental origins based on expression of the neural placode marker *Phox2b* ^35^: PSN2 through 8 and PSN-H express high levels of *Phox2b*, indicating derivation from the neural placode, whereas PSN1 and PSN9 lack detectable *Phox2b* expression and are therefore likely neural crest–derived (Figure 1I). Together, these findings show that the 10 molecularly defined PSN subtypes encompass distinct fiber types and developmental lineages.

We compared our molecular atlas with two previously published scRNA-seq datasets of vagal sensory neurons and identified corresponding clusters based on molecular marker genes presented in those studies (Figure S2A and B). In one study with nodose sensory neurons retrogradely labeled from the mouse lungs ^27^, all four major lung clusters have corresponding subtypes from our result. In addition to the two jugular subtypes, which were not the focus of the previous study, three additional nodose subtypes (PSN3, PSN6, and PSN8) were revealed here, likely due to more efficient labeling of these neurons by WGA compared to AAV instillation (Figure 1D). Overall, our molecular profiling uncovered the most comprehensive diversity of vagal PSNs to date.

### PSN subtype markers and abundance in vivo

To identify specific markers for each PSN subtype, we applied the single-cell differential expression (SCDE) method ^36^, comparing the expression profile of each subtype against every other subtype individually. We then intersected the resulting lists of differentially expressed genes, which yielded 10 to 89 genes enriched in each PSN subtype (except for the hybrid subtype PSN-H), totaling 260 genes for the 9 subtypes (Figure 2A and S2C). From these subtype-enriched gene sets, we selected the most specific and robust markers for each subtype based on their expression specificity and levels (Table S2). This list includes some genes previously identified marking subpopulations of PSNs, such as *Calb1, Npy2r,* and *Gfra3* ^17,24,37^, however, many previously used markers of mouse PSN subpopulations, such as *Trpv1*, *Piezo2*, *P2ry1*, *Slc7a7*, *P2rx2*, *P2rx3*, and *Pvalb* ^17,24,37–39^, were not among the 260 subtype-enriched genes, because they are expressed across multiple PSN subtypes (Figure S2B). Thus, our molecular atlas identified specific and robust molecular markers that allow identification, manipulation, and functional interrogation of 9 of the 10 PSN molecular subtypes we defined (all except hybrid subtype PSN-H).

To determine the *in vivo* abundance of each PSN subtype, we analyzed the expression of highly specific and robust markers using RNAscope *in situ* hybridization and/or immunostaining on vagal ganglion sections/whole-mount tissues, combined with WGA retrograde labeling of PSNs. This analysis identified all 10 PSN subtypes, including the hybrid subtype PSN-H (a subset of which are *Npy2r*⁺/*Htr3b*⁺) and PSN9 (*Phox2b*-*Gal*⁻ or *Phox2b*-*Trpv1*⁻) (Figure 2B–D). Cell counts revealed marked differences in subtype abundance, ranging from 2.5% (∼10 neurons per side) for PSN9 to 15–20% (60–80 neurons) for PSN2, PSN3, and PSN8 (Figure S2D). These proportions closely matched our scRNA-seq data, suggesting minimal cell type bias during our manual cell picking (Figure 2E). The combined abundance of all 10 subtypes accounted for nearly all WGA-labeled PSNs (Figure 2F), indicating that we obtained a near-complete molecular census of these neurons.

### PSNs in detecting physiological and pathological signals

Classical studies have shown that PSNs can be activated or modulated by a wide range of physiologically relevant stimuli, including chemical irritants, heat, mechanical deformation, and changes in ambient gas composition ^12,15,40,41^. To predict which PSN subtypes are capable of detecting these stimuli, we screened for the expression of genes known to be involved in external and internal sensing, such as mechanoreceptors, thermoreceptors, receptors for volatile and non-volatile chemicals, and their gene family members (Figure 3A and Table S3). This analysis identified subtypes associated with established PSN sensory functions described in classical studies, along with putative sensors and potential new sensory roles. Notably, it revealed that each PSN subtype likely responds to multiple types of stimuli through distinct combinations of sensors, and that most stimuli can, in turn, be detected by more than one PSN subtype.

**Figure 3.**
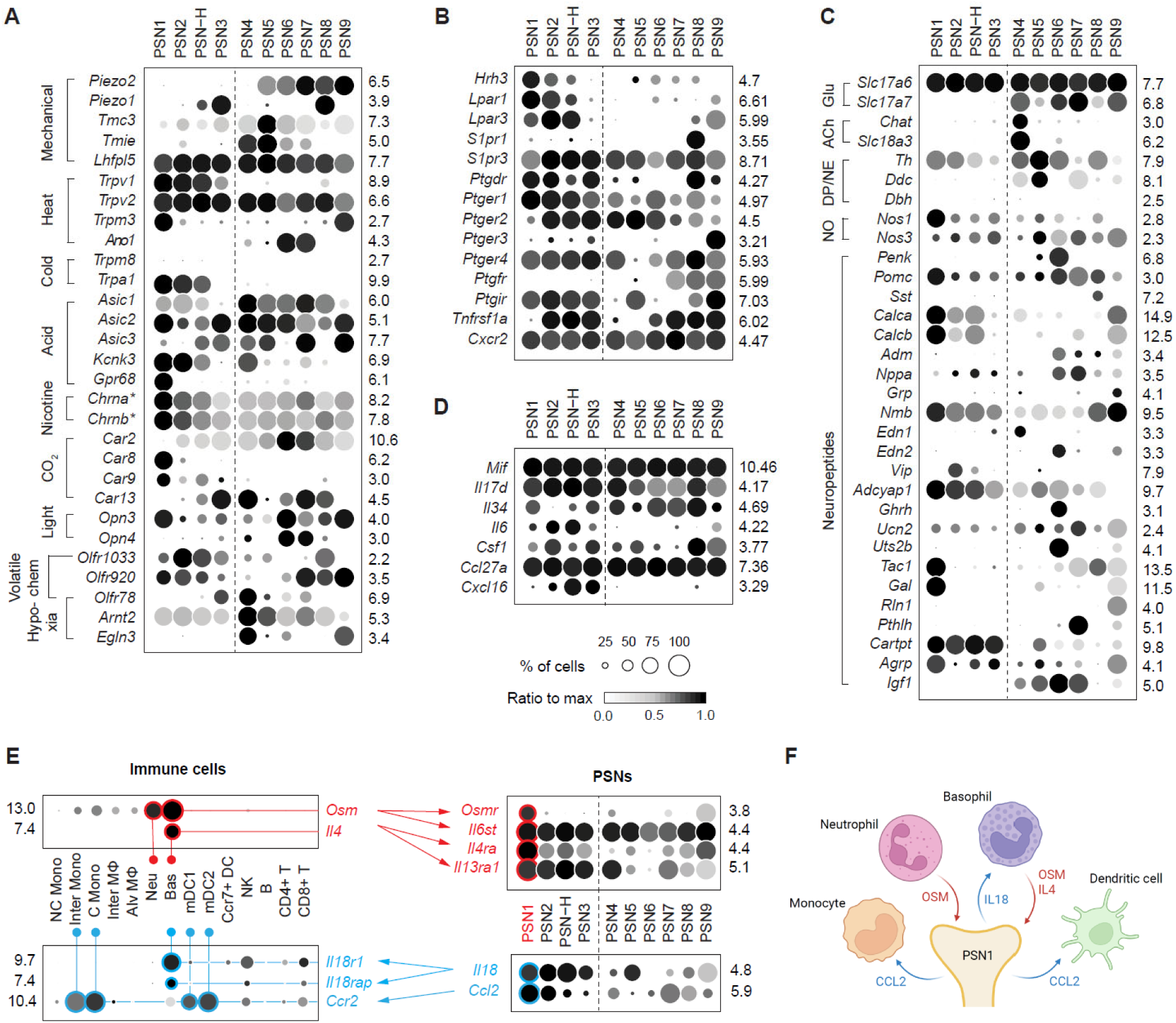
Diversity of PSNs in sensing and signaling. (A) Expression patterns of genes encoding receptors that detect diverse physiological stimuli. Expression levels of nicotinic acetylcholine receptor subunits (*) are the summed values of all isoforms (see Figure S3A for individual isoforms). (B) Expression patterns of genes encoding receptors for inflammatory mediators. (C) Expression patterns of genes required for neurotransmitter synthesis and release and genes encoding neuropeptides. Glu, glutamate; ACh, acetylcholine, DP, dopamine, NE, norepinephrine, NO, nitric oxide. (D) Expression patterns of genes encoding selected cytokines. (E) Ligand-receptor pairs expressed by lung residential immune cells and PSNs. (F) Schematics show predicted signaling interactions between lung residential immune cells and PSN1 neurons.

Hyperthermia and chemical irritants such as acids, ozone, and toxic aldehydes can stimulate pulmonary C-fibers or potentiate their responses to other stimuli ^41–45^. Many receptors detecting these stimuli are expressed by PSNs, and some of them are not limited to the Group I subtypes (e.g., thermal receptors *Trpv2* and *Trpm3*, acid-sensing channels *Asic1-3*), suggesting that these signals may also modulate the activity of A-fiber subtypes. The polymodal irritant receptor *Trpa1,* which can be activated by many chemical irritants, including aldehydes present in cigarette smoke (CS) and by cold temperature ^46,47^, is selectively expressed in PSN1 and PSN2 (and a subset of PSN-H) neurons, but not in the other Group I subtype PSN3. Hence, PSN1 and PSN2 subtypes represent classical pulmonary C-fibers that detect harmful stimuli. Interestingly, PSN1 also expresses the highest levels of nicotinic acetylcholine receptors (Figure S3A) that detect nicotine in CS and exclusively expresses the acid-sensing GPCR *Gpr68*, which has a high activation threshold (∼pH6.8) close to physiological pH ^48^, suggesting that PSN1 may be more sensitive to those irritants than PSN2.

Six of the 10 subtypes (plus a subset of PSN-H neurons) express mechanosensitive Piezo channels, indicating they are putative mechanoreceptors. These include all Group II subtypes (except PSN4) expressing *Piezo2*, and the *Trpa1*^-^ Group I subtype PSN3 (and a subset of PSN-H neurons) expressing *Piezo1;* PSN8 is the only subtype that co-expresses both Piezo channels. PSN5, in addition to *Piezo2*, also expresses *Tmc3*, a family member of the auditory hair cell mechanotransduction channels *Tmc1/2* ^49^, along with the other two components of the mechanotransduction complex, *Tmie* and *Lhfpl5* ^50,51^. The different mechanosensors and their combinations expressed by individual subtypes may contribute to their detection of different types of mechanical stimuli, such as inflation vs. lung compliance, or to their different response thresholds. Additional polymodal mechanosensitive channels, such as two-pore potassium channels and acid-sensing ion channels, are also expressed in multiple subtypes, including PSN1 and PSN2, suggesting that these subtypes may also be modulated by mechanical stimuli.

Hypercapnia (high CO_2_) has been shown to increase the activity of pulmonary C-fibers and suppress the activity of pulmonary mechanoreceptors, the latter depending on carbonic anhydrase (CA) ^52,53^. Multiple CA genes are expressed by PSNs, and several (*Car2*, *Car8*, *Car9, Car13)* are selectively enriched in subsets of PSN subtypes. *Car2*, which mediates olfactory detection of CO_2_ ^54^, has the highest expression levels and is selective for two putative mechanosensory subtypes (PSN6 and 7). As we show below, these two subtypes control eupneic breathing, suggesting that the control mechanisms may be inactivated under hypercapnia by *Car2*-mediated suppression of their mechanosensory functions.

We also uncovered the expression of genes that mediate sensory functions previously unknown in PSNs, including light-sensitive non-visual opsins *Opn3* and *Opn4,* the latter selectively expressed in PSN6 and 7. Several olfactory and vomeronasal receptors, such as *Olfr1033*, *Olfr78,* and *Olfr920*, were also detected, and they are differentially expressed across PSN subtypes. Among them, *Olfr78*, enriched in PSN3 and 4, is of particular interest because it has been shown to respond to lactate and mediate the acute hypoxia response in carotid bodies ^55^. PSN4 also expresses *Kcnk3*/TASK-1, another acute hypoxia response mediator, and high levels of genes in the HIF chronic hypoxia response pathway *(Arnt2/*Hif2b, *Egln3*/Phd3) ^56^. Together, these suggest that PSN4 may be involved in hypoxia detection or adaptation. We also found that PSNs express receptors for cellular components specific to bacteria (*Tlr4*, *Tlr5*, *Nod1*, and *Naip1*), cell surface proteins that enable entry of respiratory viruses (e.g., Uvrag for influenza, Igf1r for respiratory syncytial virus (RSV), and Pvrl4 for measles), and intracellular RNA and DNA sensors (*Ddx58*, *Dhx58*, *Ddx41*, *Lrrfip1* and *Aim2*), suggesting they can direct detect or be influenced by pathogens (Figure S3B and S3C).

In addition to detecting physiological stimuli, PSNs express a remarkable variety of receptors for inflammatory signals including purines (e.g., ATP and adenosine), biogenic amines (serotonin and histamine), lipids (e.g., lysophosphatidic acid, sphingosine-1-phosphate, and prostaglandins), proteases (e.g., thrombin and trypsin) and cytokines (e.g., interleukins, chemokines and tumor necrosis factors) (Figures 3C, S3D and S4). For many inflammatory signals, the receptors are expressed across all PSN subtypes (e.g., ATP and prostaglandins receptors and *Cxcr2*), although some with different combinations of receptor isoforms. This implies widespread influence of inflammation on PSNs. However, other signal receptors enriched in specific subtypes (e.g., lysophosphatidic acid receptors in PSN1, 2, and H and TNF-α receptor in PSN2, H, 3, 7, 8, and 9) or even in a single subtype (e.g., oncostatin M receptor and interleukin 4 receptor in PSN1). Thus, each PSN subtype appears to detect or be modulated by multiple, overlapping yet distinct combinations of inflammatory signals, as for their detection of physiological signals.

### PSNs in secreting signaling molecules and interacting with lung immune cells

As pseudounipolar neurons, PSNs communicate with both cells in the lungs and secondary neurons in the brainstem. To identify the neuronal signals used by each PSN subtype, we first analyzed expression of genes encoding neurotransmitter synthesis and secretion machinery and neuropeptides. As expected from previous studies, all are glutamatergic (all express *Slc17a6* and a subset *Slc17a7*), but we also identified two novel PSN neurotransmitters: acetylcholine in PSN4 (expresses *Chat* and *Slc18a3*) and dopamine in PSN5 (expresses *Th* and *Ddc*, but not *Dbh*). We also found that nitric oxide synthase genes (*Nos1* and *Nos3*) are widely expressed across PSN subtypes, with PSN1 expressing neuronal NOS (*Nos1*) at the highest level, suggesting the use of NO. A surprisingly large number of neuropeptide genes (77%, 57 out of 74 known genes) were detected in PSNs (Table S4) and 23 of these were expressed by considerable number of PSNs (in >50% PSNs in at least one subtype) and exhibited subtype-selective expression patterns including several that are specific for a single subtype (*Gal* in PSN1, *Penk*, *Uts2b* and *Ghrh* in PSN6, and *Pthlh* in PSN7) (Figure 3C).

Given the emerging importance of neuro-immune interactions in maintaining lung homeostasis and responding to diseases ^57,58^, in addition to neurotransmitters and neuropeptides, we further explored the potential PSN-derived signals that influence immune cells by screen through genes encoding cytokines and chemokines. Surprisingly, we identified dozens of these genes expressed by PSNs, with some expressed across all PSN subtypes (e.g. *Mif* and *Ccl27a*), and some enriched in subsets of subtypes (e.g., *Il6* and *Cxcl16*) (Figure 3D). To further determine if there are potential interactions between PSNs and immune cells in basal condition or readily upon a challenge, we analyzed a previously published mouse lung cell atlas ^59,60^ for expression of receptors of these PSN-expressing cytokines and chemokines. We found these receptors are differentially expressed across lung immune cell types (e.g., *Csf1r* mainly in monocytes and interstitial macrophages and *Cxcr6* in CD8^+^ T cells), and collectively they are distributed across all types of immune cells, including both innate and adaptive immune cells (Figure S4). Given that PSNs also express receptors for cytokines and chemokines (Figure 3B, 3E and S4), some of which are found expressed by lung immune cells, this bidirectional cytokine signaling between PSNs and immune cells (Figure 3E, 3F and S4) suggests that PSNs potentially bridge different types of immune cells in an immune response cascade or amplify the response in certain immune cells.

### Group I PSN subtypes form free nerve endings with terminals in different lung compartments

The lung is composed of a serial branching bronchial tree (the conducting airways) and millions of alveoli, where gas exchange occurs, together with an intricate vascular network. To begin to define the receptive fields and sensory mechanisms of PSN subtypes, we devised genetic labeling strategies with subtype selective genes to map subtype terminal distributions, morphologies, and cell contacts in the lung. Subtype labeling was performed by injecting an adeno-associated virus (AAV) expressing a Cre-dependent reporter (fluorescent protein or alkaline phosphatase, AP) into the vagal ganglia (or instilling them into the lungs) of mice expressing Cre recombinase driven by a gene selectively expressed by the subtype(s) of interest (Figure 4A).

**Figure 4.**
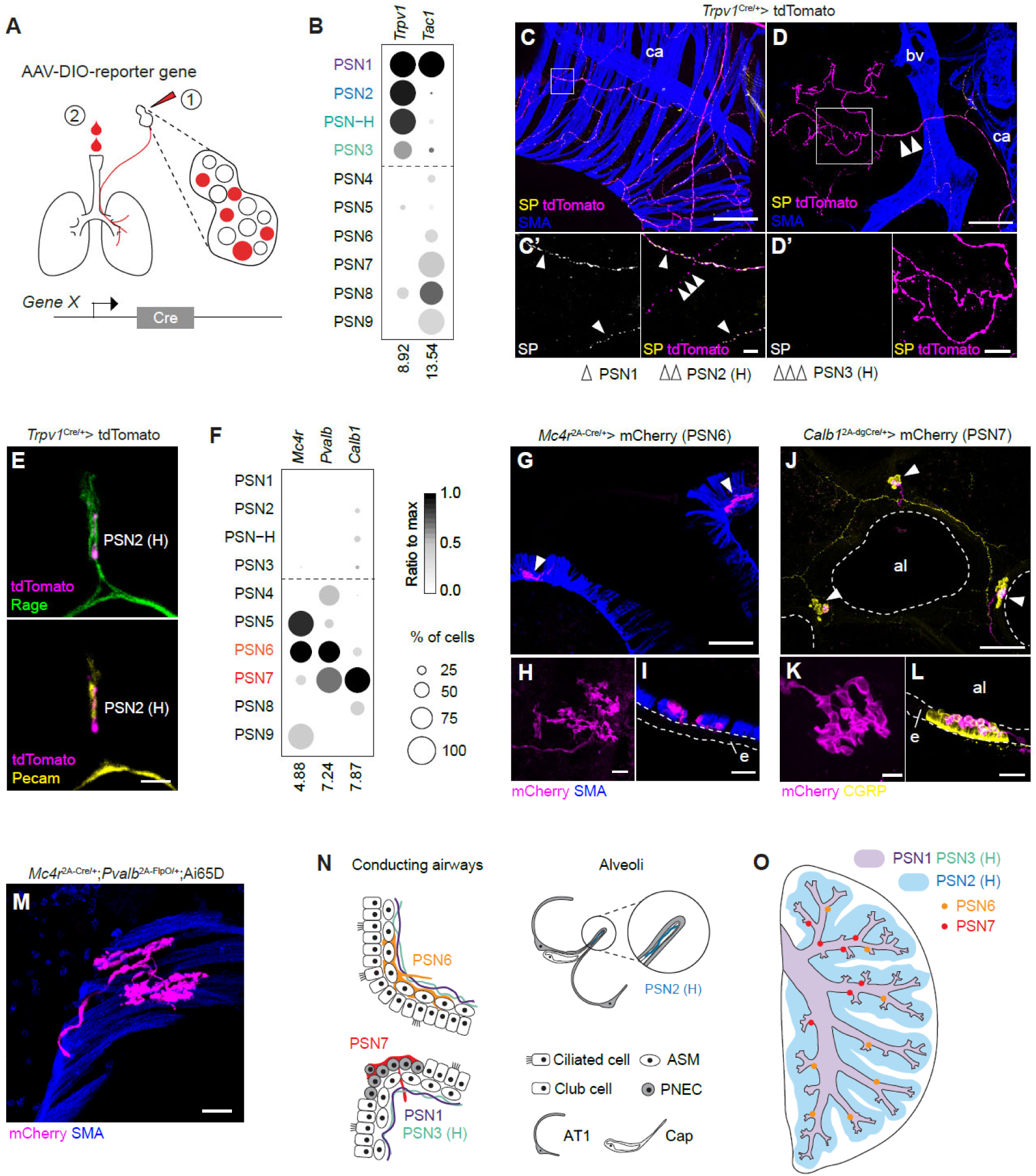
Diversity of PSNs in terminal morphologies and locations. (A) Labeling strategy for PSN subtype neurons. AAVs expressing Cre-dependent reporter genes (fluorescent proteins or alkaline phosphatase) were injected into the vagal ganglia (1) or instilled into the lungs (2) of mice expressing Cre recombinase driven by a gene (promoter of gene X in diagram) selectively expressed by the subtype(s) of interest. (B) Expression of *Tac1* differentiates PSN1 from PSN2, H, and 3 among *Trpv1*-expressing PSNs.(C-D) Group I neurons labeled by injecting AAV-CAG-FLEX-tdTomato into the vagal ganglia of *Trpv1*^Cre/+^ mice. C and D, maximum projections of a segment of conducting airways (C) and a region mainly occupied by alveoli (D). Sections were immunostained for substance P (SP, encoded by *Tac1*), tdTomato and smooth muscle actin (SMA, labels smooth muscle cells that surround conducting airways and major blood vessels and are absent in alveolar regions). Close ups of boxed regions are shown in C’ and D’. C’ shows both SP+ (single arrowheads, PSN1) and SP-(triple arrowhead, likely PSN3 or H) labeled fibers. D’ shows a SP-labeled fiber (double arrowhead, likely PSN2 or H) innervating alveoli. Note that virtually all alveolar tdTomato+ fibers were SP-. bv, blood vessel; ca, conducting airway. Scale bars: 100 μm (C and D), 10 μm (C’), 20 μm (D’) (E) A thin optical section of alveoli immunostained with tdTomato, Rage (alveolar type I cell marker), and Pecam (capillary marker), showing an alveolar Group I neuron terminal, presumably a PSN2 or H neuron, contacting both alveolar cell types. Scale bar: 5 μm. (F) Expression patterns of genes selectively expressed by PSN6 and PSN7 neurons. (G-I) PSN6 terminals labeled by injecting AAV-Syn-DIO-hM3Dq-mCherry into the vagal ganglia of *Mc4r*^2A-Cre/+^ mice. G, maximum projection of a bronchial branch point with two labeled terminals (arrowheads). H, top-down view of a labeled terminal showing multi-branched “leaf” shape morphology. I, side view of a labeled terminal intercalating with airway smooth muscle fibers (SMA+). al, airway lumen; e, epithelium (outlined). Scale bars: 100 μm (G), 20 μm (H and I). (J-L) PSN7 terminals labeled by injecting AAV-Syn-DIO-hM3Dq-mCherry into the vagal ganglia of *Calb1*^2A-dgCre/+^ mice, followed by TMP treatment. J, maximum projection of conducting airways with three NEBs (labeled by CGRP immunostaining) all innervated by labeled terminals (arrowheads). K, top-down view of a labeled terminal showing “waffle” shape morphology, L, side view of a labeled terminal covering the apical side of the NEB. al, airway lumen; e, epithelium (outlined). Scale bars: 100 μm (J), 10μm (K), 20μm (L). (M) A PSN6 terminal labeled by tdTomato in a *Mc4r*^2A-Cre/+^;*Pvalb*^2A-FlpO/+^;Ai65D mouse. Only PSN6 neurons are double positive for *Mc4r* and *Pvalb*. Scale bar: 20 μm. (N) Diagrams of 6 PSN subtypes showing their terminal structures, locations, and contact lung cells. PSN1, 2, H and 3 all form free nerve endings. PSN1 and 3 both branches along conducting airways beneath airway smooth muscle. PSN2 mainly branches along the alveolar junctions and closely contacts alveolar type I and capillary cells. PSN6 terminals intercalate with airway smooth muscle, preferentially at bronchial branch points. PSN7 also terminates at branch points but on the apical side of NEBs. ASM, airway smooth muscle; PNEC, pulmonary neuroendocrine cell; AT1, alveolar type I cell; Cap, capillary cell. (O) Distribution of PSN subtypes throughout a lung lobe. PSN1 and PSN3 both terminate on conducting airways (light purple), whereas PSN2 terminates predominately in the alveolar region (light blue) (whether PSN3 also innervates alveoli is undetermined). PSN-H likely exhibits a similar distribution as PSN2 or PSN3 or a combination of both. Group I neurons also terminate on pulmonary blood vessels (not shown). PSN6 (orange dots) and PSN7 (red dots) both terminate at bronchial branch points but with different proximal and distal distributions as shown. However, the terminal locations of PSN6 and 7 neurons are not stereotyped between different lobes or animals, and the schematic only illustrates general features of their terminal distributions.

Terminals of Group I neurons were labeled by injecting AAV-CAG-DIO-tdTomato into the vagal ganglia of *Trpv1*^Cre/+^ mice, since *Trpv1* is expressed in the vast majority of Group I neurons but rarely in Group II neurons (Figure 4B). TdTomato-labeled axons exhibited simple terminal structures (free-nerve endings) and terminated throughout the bronchial tree and alveoli (Figure 4C and 4D). To distinguish axons of PSN1 neurons from other Group I neurons, we co-stained lung sections with an antibody against substance P (SP), a neuropeptide encoded by *Tac1*, which is selectively expressed by PSN1 among Group I neurons (Figure 4B). TdTomato^+^SP^+^ fibers ramified beneath the airway smooth muscle fibers throughout the conducting airways, with limited penetration of the alveolar region (Figure 4C, 4D and S5A). These fibers also terminated on blood vessels, generally external to the vascular smooth muscle layer (Figure S5E and S5F). Thus, the sensory field of PSN1 neurons is the peribronchial and perivascular mesenchyme. Tdtomato^+^SP^-^ fibers, presumably from PSN2, H and 3 neurons, terminate along the conducting airways (Figure 4C), on blood vessels (Figure S5G), and in the alveolar region (Figure 4D). In the alveolar region, these fibers contact both epithelial and capillary endothelial cells (Figure 4E), and they branch extensively, running mainly along the boundaries between alveoli, with rare cases that they elaborate on air sacs (Figure S5B). PSN2 neurons (and likely some PSN-H neurons) almost exclusively form alveolar terminals and only rarely terminate on bronchial airways, as shown by whole-mount alkaline phosphatase (AP) staining of lungs from *Npy2r*^ires-Cre/+^ mice intratracheally instilled with AAV expressing Cre-dependent AP (Figure S5C and S5D). This result is consistent with a prior study that injected a reporter virus into the vagal ganglia of *Npy2r*^ires-Cre/+^ mice ^24^. Thus, the major sensory field of PSN2 neurons is the alveoli, and the tdTomato^+^SP^-^ fibers that terminated on conducting airways (observed in *Trpv1*^Cre/+^ mice) most likely emanated from PSN3 (and some PSN-H) neurons (Figure 4C, 4N and 4O).

### PSN6 and PSN7 neurons terminate on airway smooth muscle and neuroepithelial bodies respectively

We also mapped the terminals of two Group II subtypes, PSN6 and PSN7, which we subsequently functionally characterized (see below). We labeled PSN6 neurons by injecting either AAV-CAG-DIO-tdTomato or AAV-Syn-DIO-hM3Dq-mCherry into the vagal ganglia of *Mc4r*^2A-Cre/+^ mice (the latter AAV labeled membrane structures so better revealed terminal morphologies). Although *Mc4r* is also expressed in PSN5 neurons (Figure 5F and S5H), we found that AAV infection in the ganglion preferentially labeled PSN6 neurons, potentially due to viral tropism (Figure S5I and S5J). We found that mCherry labeling in the lung was, with rare exception, confined to multi-branched, "leaf" shape terminals intercalating between airway smooth muscle fibers beneath the airway epithelium (Figure 4H and 4I). These resemble the smooth muscle-associated terminals observed in humans and other species ^61,62^. Interestingly, PSN6 terminals were not uniformly distributed across airway smooth muscle: a large majority of terminals localized to smooth muscle at bronchial branch points (Figure 4G and S5K), and whole-mount staining of AP-labeled PSN6 fibers showed them terminating mainly on primary and secondary bronchial branches and concentrated centrally in lobes (Figure S5N and S3Q). To further confirm the terminal location and morphology, we generated *Mc4r^2A-Cre/+^*; *Pvalb^2A-FlpO/+^*; Ai65/+ triple heterozygous mice, in which only *Mc4r* and *Pvalb* double positive neurons were labeled, excluding PSN5 neurons (*Pvalb*^-^) (Figure 4F). In these mice, we observed tdTomato-labeled terminals with the morphology described above and they are only present at the airway branch points (Figure 4M). We conclude that the PSN6 sensory field is airway smooth muscle cells at branch points, mainly in the middle region of the bronchial tree (Figure 4N and 4O). It appeared that most of the individually labeled fibers form a small number of terminals (1∼4) within close proximity (Figure S5R), suggesting a restricted receptive field for individual neurons.

**Figure 5.**
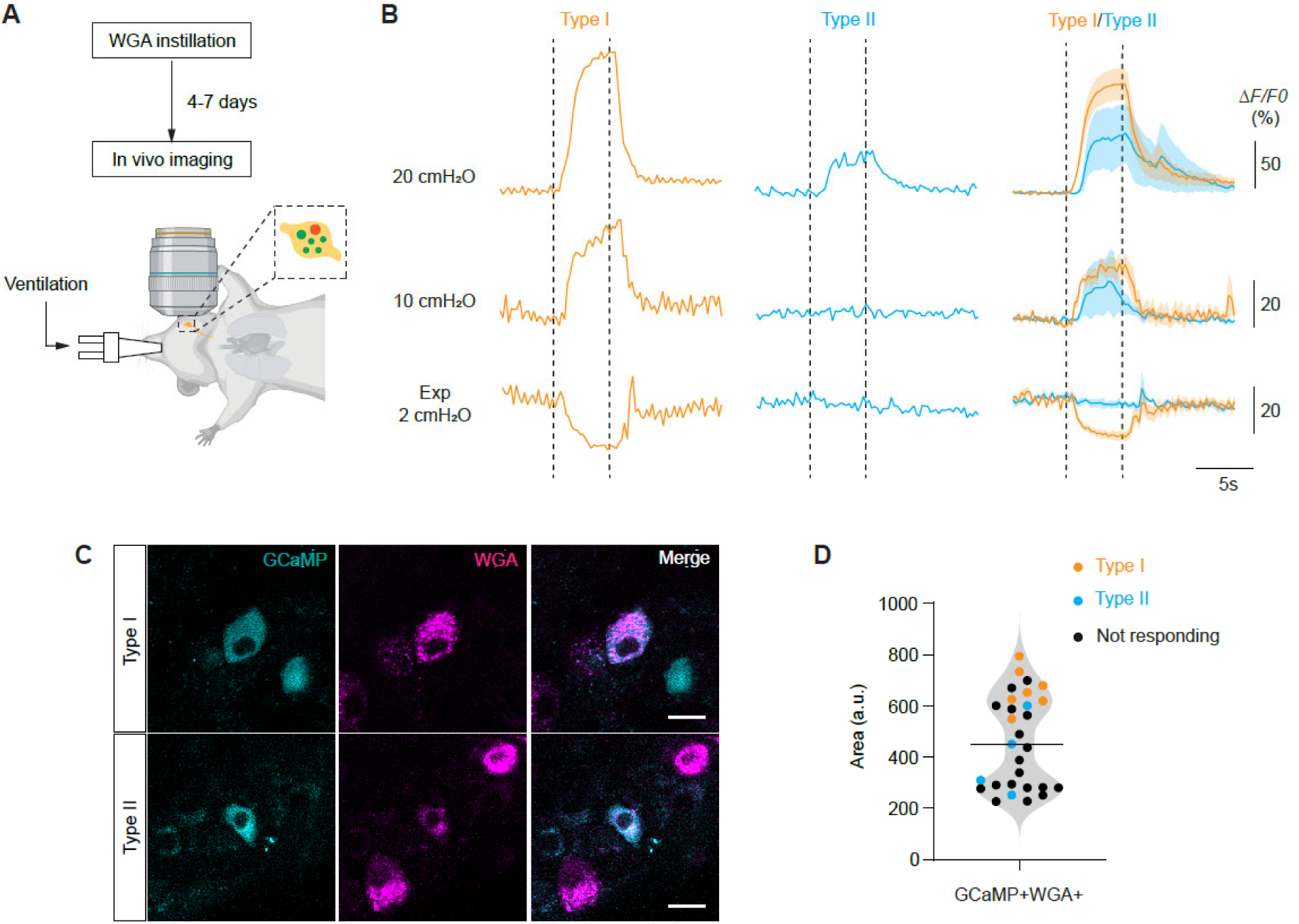
PSN6 neurons are activated by lung inflation. (A) Schematic of the *in vivo* calcium imaging workflow. WGA-594 was instilled prior to imaging experiments to label PSNs. Imaging was performed on anesthetized, intubated, and mechanically ventilated mice. (B) Representative and averaged traces of neuronal responses to 5s sustained lung inflation (20 cmH_2_O and 10 cmH_2_O) and deflation (2 cmH_2_O). Dashed lines depict the start and end of the holding period. Neurons were classified into two response types based on whether their GCaMP signals decreased during deflation: Type I (orange), decreased; Type II (cyan), no decrease. (C) Example images of neurons corresponding to the two response types shown in panel B. (D) Cell body size distributions of recorded neurons (29 neurons from 4 mice). Neurons were identified in Z-stacks of the vagal ganglion acquired prior to time-series recordings, and the optical plane with the largest cross-sectional area was used for quantification.

PSN7 neurons were labeled by injecting AAV-Syn-DIO-hM3Dq-mCherry into the vagal ganglia of *Calb1*^2A-dgCre/+^ mice, in which any expressed destabilized Cre protein (dgCre) is rapidly degraded unless stabilized by trimethoprim (TMP) ^63^. TMP delivery (300µg/g body weight) two days after viral injection specifically labeled Calb1^+^ neurons among WGA-labeled PSNs (Figure S5L and S5M). All mCherry^+^ fibers terminated on neuroepithelial bodies (NEBs) (Figure 4J), clusters of neurosensory epithelial cells (neuroendocrine cells) located at bronchial branch points ^64^, where Calb1-stained neurites have previously been observed in immunostained lung sections ^37^. Labeled PSN7 axons penetrated the basement membrane of the epithelial layer of the NEBs, coursed between neuroendocrine cells, and terminated in an elaborate "waffle" shape covering their apical surface, facing the airway lumen (Figure 4K and 4L). Whole-mount staining of AP-labeled PSN7 axons showed that they also terminated on major bronchial branches and were enriched in the proximal regions of the lobes (Figure S5O and S5Q). We conclude that the PSN7 sensory field is the airway surface of neuroendocrine cells at bronchial branch points, mainly in the proximal region of the bronchial tree (Figure 5N and 5O). A substantial fraction of labeled fibers forms multiple terminals, which can be distributed through multiple branch points (Figure S5S), suggesting a broader receptive field than PSN6 neurons.

Thus, molecularly distinct subtypes of both Group I and Group II PSN subtypes have different lung receptive fields, and their terminals have different morphologies and target cells.

### PSN6 neurons detect lung inflation

The close association with airway smooth muscle fibers and its expression of mechanosensitive channel *Piezo2* suggest that PSN6 neurons detect smooth muscle stretch during lung inflation. To assess the response properties of these neurons to lung inflation, we crossed *Mc4r^2A-Cre/+^* line with Cre-dependent GCaMP lines (Ai148 or Ai95) and performed *in vivo* calcium imaging on the vagal ganglion (Figure 5A). We used a mechanical ventilator to ventilate the mice with parameters that closely recapitulate eupneic breathing patterns (150 breath/minute, 10-12 cmH2O target pressure, I/E ratio=1:2). Static holding after inspiration (two different levels: 10 and 20 cmH_2_O) or expiration (at 2 cmH_2_O) were introduced. To identify which GCaMP6f expressing neurons innervate the lungs, we instilled fluorescent conjugated WGA 4-7 days prior to the imaging session and selected WGA and GCaMP double positive cells for imaging. Among these neurons, we found a subset (11 out of 29 neurons from four mice) increasing their Ca^2+^ signal intensity upon inflation holding (at 20 cmH_2_O) and the increase sustained the holding period, indicating these responding neurons are slowly-adapting mechanoreceptors (Figure 5B). Among the 11 neurons responding to inflation holding at 20 cmH_2_O, 7 had increased Ca^2+^ signal levels at 10 cmH_2_O and decreased levels during expiration holding (Type I neurons), indicating that these neurons have a low mechanical threshold and are active during normal ventilation cycles (Figure 5B). The other four neurons responding to 20 cmH_2_O inflation did not decrease their Ca^2+^ signal intensity during expiration holding (Type II), suggesting these neurons are likely not active during normal ventilation cycles. Both PSN6 and PSN5 neurons are *Mc4r*^+^, therefore they are both recorded in the experiment. Given that PSN6 have larger cell bodies than PSN5 neurons (Figure 1G), we quantified cell body sizes of imaged neurons (Figure 5C and 5D). We found that all neurons with Type I response profile are among the large diameter neurons, therefore PSN6 neurons. Thus, we conclude that PSN6 neurons, at least a subset if not all, are slowly-adapting low-threshold inflation receptors, and their activity reports the inflation level of the lungs during each inspiration.

### PSN6 neurons regulate inspiratory time

To further explore the functions of PSN6 neurons in regulating breathing, we selectively eliminated them using the diphtheria toxin (DT)/ DT receptor (DTR) cell ablation system. DTR was expressed in PSN6 neurons by bilaterally injecting AAV-CBA-DIO-DTR-GFP, which expresses a Cre-dependent DTR-GFP fusion transgene ^65^, into the vagal ganglia of mice carrying the PSN6-selective Cre-driver *Mc4r*^2A-Cre^ (PSN6-DTR mice). Three weeks after viral injection, DTR-GFP was detected in 35-55% of PSN6 neurons (*Lmcd1*^+^WGA^+^) (Figure S6A and S6B), and three days after subsequent systemic delivery of DT to induce apoptosis of DTR-GFP expressing cells, no DTR-GFP expressing PSNs were detected in the ganglia (Figure S6C and S6D). To assess the impact of PSN6 loss on breathing, we first performed whole-body plethysmography recordings on freely moving control (wildtype mice injected with the same virus) and PSN6-DTR mice 1 day before and 3 days after DT injection (Figure 6A), focusing the analysis on eupneic ("quiet") breaths computationally selected based on their regular breathing pattern (Figure S6E and S6F). We found that after DT injection the tidal volume (TV) of eupneic breaths in PSN6-DTR mice was increased by ∼11% over control mice (Figure 6B and 6C). Inspiratory time (Ti) was extended to a similar degree (Figure 6D-F), accounting for the entire increase in tidal volume. The effect was specific for inspiratory time because inspiratory flow, the other determinant of tidal volume, was not significantly altered (Figure 6G). Expiratory time (Te), Peak expiratory flow (PEF), respiratory frequency (f), and minute ventilation (MV) did not show significant changes (Figure S6G-J). These results indicate that PSN6 neurons are required for setting the eupneic tidal volume, and they appear to do so by regulating inspiratory time.

**Figure 6.**
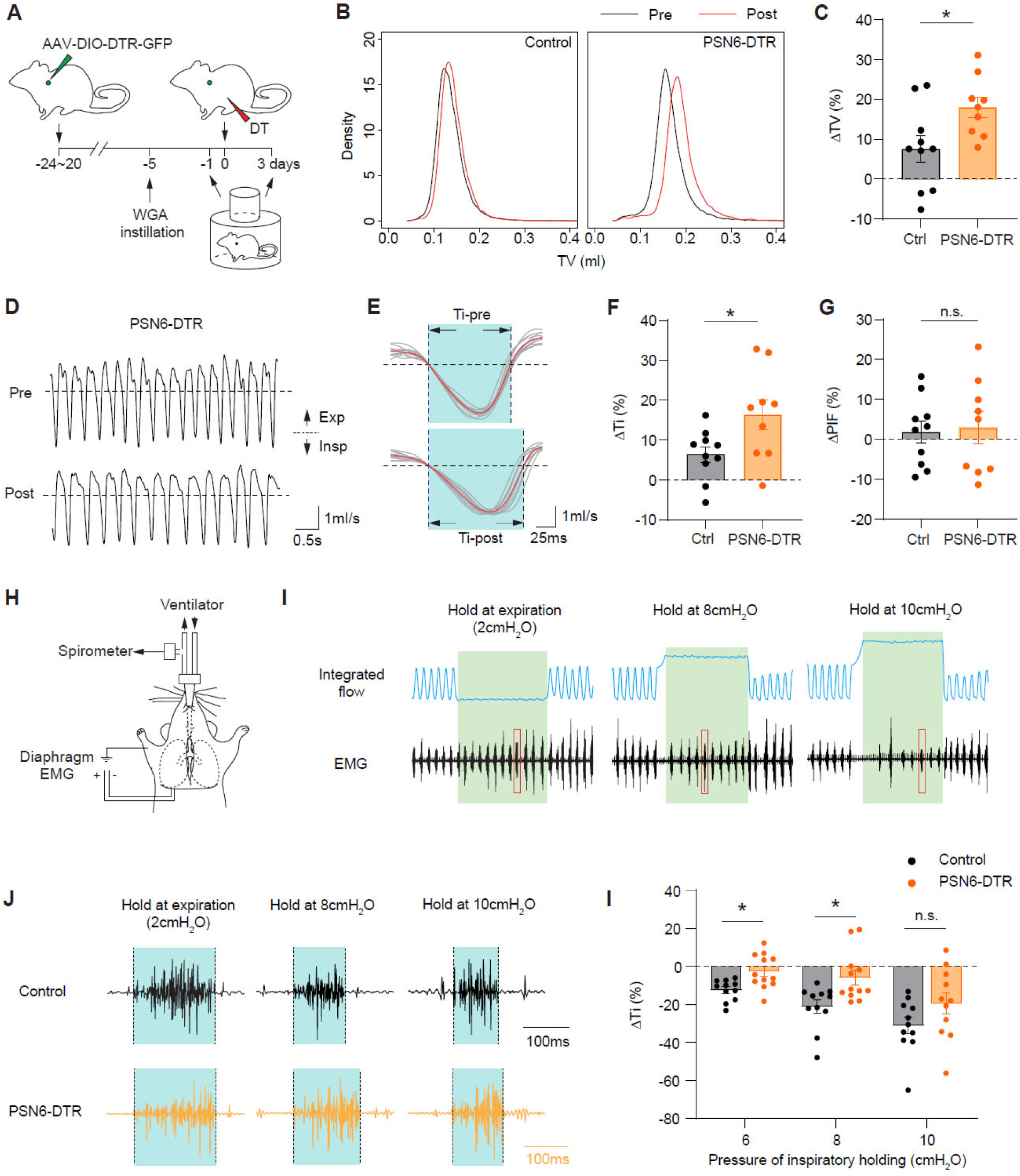
Ablating PSN6 neurons increases tidal volume and inspiratory time. (A) Experimental scheme for assessing functions of PSN subtypes in breathing regulation. AAV-DIO-DTR-GFP was bilaterally injected into the vagal ganglia of subtype-specific Cre mice. Three weeks (20-24 days) later, DT was injected intraperitoneally to ablate the targeted PSN subtypes. Whole-body plethysmography (WBP) recordings to acquire breathing parameters were performed 1 day before (pre-ablation) and 3 days after DT injection (post-ablation). WGA retrograde labeling was performed 5 days prior to DT injection to label PSNs for later assessment of ablation efficiency. (B) Kernel density plots showing tidal volume (TV) distribution of calm breaths of a control and a PSN6-DTR mouse pre- and post-DT injection. Note shift to higher tidal volumes following PSN6 ablation. (C) Tidal volume (TV) changes after PSN6 ablation. Each dot represents the post-DT change in the TV distribution peak, normalized to the pre-DT peak value. Bars, mean ΔTV. Error bars, standard error of the mean (s.e.m.). *p=0.028, two-sided Mann-Whitney U test. (D) Representative breathing airflow traces of a PSN6-DTR mouse before and after DT injection. Note that in the same period, there are 16 breaths before injection and 13 breaths after injection. Insp, inspiration; Exp, expiration. Dashed lines, zero flow. (E) Overlay of 10 breaths from each of the traces in D, aligned to the onset of inspiration. Red traces, average of the 10 breaths; light blue, inspiratory phase. Note the lengthened inspiratory time (Ti) after PSN6 ablation. (F, G) Inspiratory time (Ti) and peak inspiratory flow (PIF) changes after PSN6 ablation. Each dot represents the post-DT change in the Ti or PIF distribution peak, normalized to the pre-DT peak value. Note increase in Ti (F) but not PIF (G) following PSN6 ablation. Error bars, error of the mean (s.e.m.). *p=0.035, two-sided Mann-Whitney U tests. (H) Inflation challenge assay. Open-chest mice were intubated and ventilated. Airflow was monitored by spirometry, and inspiratory drive was measured by electromyogram (EMG) recording of the costal diaphragm. (I) Representative traces of integrated airflow (i.e., tidal volume) and diaphragm EMG from a control mouse during sustained expiratory (at 2cm H_2_O) and inspiratory (at 8cm and 10cm H_2_O) holdings (5s each, green shadings). Note decrease in the rate of EMG bursts along with increase in the pressure maintained during inspiratory holdings. Red boxes, single bursts of EMG activity enlarged in (J). (J) Representative traces of single diagram EMG activity bursts from control and PSN6-DTR mice after DT injections. Note longer inspiratory time (period of a single EMG burst) at 8cm H_2_O in PSN6 ablated mouse than in control. (K) Quantification of changes in Ti in control and PSN6-DTR mice after DT injections. ΔTi represents the change in the average length of EMG bursts during the first 2 seconds of inspiratory hold at a given pressure, normalized to the average length of EMG bursts during the first 2 seconds of expiratory hold. *p=0.0146 (6 cmH_2_O) and 0.0216 (8 cmH_2_O), two-way ANOVA with sidak multiple comparison correction.

*Mc4r* is not only expressed by PSNs, but also by neurons innervating other organs ^66^, therefore, we sought to further test the function of PSN6 neurons in mediating breathing reflexes to mechanical challenges directly introduced to the lungs. Given that PSN6 neurons are active during normal ventilation and respond to sustained lung inflation holding, we adopted an inflation challenge assay in which urethane-anesthetized, open-chest mice were mechanically ventilated and inspiratory motor activity was measured by concurrent electromyogram (EMG) recording of the diaphragm, an essential inspiratory muscle (Figure 6H). We introduced sustained inflations at several defined target pressures (6-20 cmH_2_O) and sustained deflations at positive end-expiratory pressure (PEEP, 2 cmH_2_O) and assessed the EMG changes upon those mechanical challenges. In control mice, we observed that inspiratory time declined and expiratory time prolonged along with the target pressure of inflation holding increased, eventually ending up with a complete apnea during the holding period. In PSN6 ablated mice, the pressure-dependent prolongation of expiration time was largely normal (Figure S6K and S6L), but the effect of inflation pressure on inspiratory time was diminished. This result is consistent with the observation of Ti prolongation in the WBP experiment, indicating that PSN6 neurons, not the *Mc4r*^+^ neurons innervating other organs, regulate inspiratory time. Interestingly, the diminution was only evident during inspiratory holding at 6 and 8 cmH_2_O, but not at higher levels of inflation (Figure 6J and 6K). We conclude that PSN6 neurons function during eupneic breathing to specifically regulate tidal volume, and they do so by detecting lung expansion with a low threshold and limiting inspiratory time.

### PSN7 neurons appear to regulate inspiratory flow

PSN7 neurons are also large diameter neurons expressing mechanoreceptor *Piezo2*, and they terminate on PNECs at the airway branch points. To assess the activity patterns of these neurons, we performed *in vivo* calcium imaging using *Calb1*^2A-dgCre/+^; Ai95/+ or *Calb1*^2A-dgCre/+^; Ai148/+ mice. We found that a small fraction of PSN7 neurons (4 out of 19 neurons from five mice) respond to high-level lung inflation (20 cmH_2_O) but not to low-level inflation (10 cmH_2_O), and their responses exhibited a rapidly-adapting property (Figure 7A and 7B). We also observed other types of responses, such as decreasing signal intensity during inflation holding (2 out of 19 neurons) (Figure 7A), suggesting that the activities of these neurons are not purely associated with lung inflation or deflation states.

**Figure 7.**
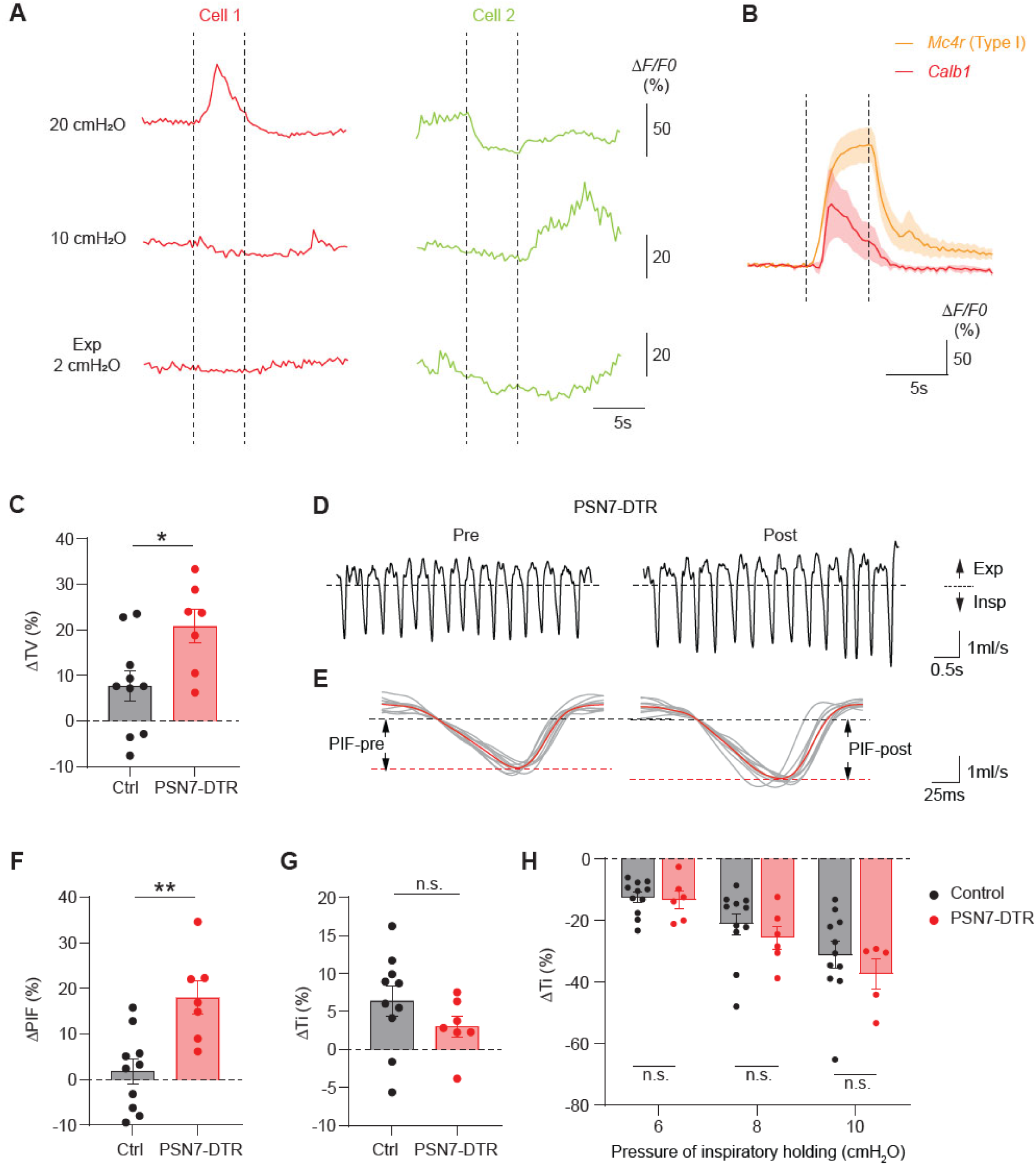
PSN7 neurons regulate inspiratory flow. (A) Representative traces of PSN7 neuronal responses to 5s sustained lung inflation (20 cmH_2_O and 10 cmH_2_O) and deflation (2 cmH_2_O). Dashed lines depict the start and end of the holding period. (B) Comparison of response properties to 20 cmH_2_O inflation holding between type I neurons among *Mc4r*-lineage PSNs and neurons with increased activity during holding among *Calb1*^+^ PSNs. Note that *Calb1*^+^ neurons exhibit rapidly-adapting property. (C) Tidal volume (TV) changes after PSN7 ablation. Each dot represents the post-DT change in the TV distribution peak, normalized to the pre-DT peak value. Bars, mean ΔTV. Error bars, standard error of the mean (s.e.m.). *p=0.043, two-sided Mann-Whitney U test. (D) Representative breathing flow traces of a PSN7-DTR mouse before and after DT injection. Insp, inspiration; Exp, expiration. Dashed line, zero flow. (E) Overlay of 10 breaths from each of the traces in B, aligned to the onset of inspiration. Red traces, average of the 10 breaths. Black dashed line, zero flow; red dashed lines, average peak inspiratory flow. Note increase in peak inspiratory flow (PIF) after PSN7 ablation (right panel). (F, G) Inspiratory time (Ti) and peak inspiratory flow (PIF) changes after PSN7 ablation. Each dot represents the post-DT change in the Ti or PIF distribution peak, normalized to the pre-DT peak value. Note increase in PIF (F) but not Ti (G) following PSN7 ablation. Error bars, error of the mean (s.e.m.). **p=0.008, two-sided Mann-Whitney U tests. Note increase in PIF (E) but not Ti (E) following PSN7 ablation. (H) Quantification of changes in Ti in control and PSN7-DTR mice after DT injections. Two-way ANOVA with Sidak multiple comparison correction.

To probe the physiological function of PSN7 neurons, we selectively ablated these neurons by injecting AAV-CMV-DIO-DTR-GFP into the vagal ganglia of PSN7-specific *Calb1*^2A-dgCre/+^ mice (PSN7-DTR mice). Three weeks after viral infection and TMP treatment, DTR-GFP was detected in 55-70% of PSN7 neurons (Calb1^+^WGA^+^) (Figure S7A and S7B), and three days after subsequent systemic delivery of DT, no DTR-GFP expressing PSNs were detected in the ganglia, with a single exception (Figure S7C and S7D). Interestingly, plethysmography recordings showed that PSN7 ablation also increased tidal volume of eupneic breaths by ∼15% over control mice, similar to the effect of PSN6 ablation described above (Figure 7C). However, instead of an increase in inspiratory time as we observed after PSN6 ablation, PSN7 ablation increased peak inspiratory flow by ∼13% over control mice (Figure 7D-G). The effect was specific as peak expiratory flow was not significantly altered (Figure S7E). Expiratory time, respiratory frequency, and minute ventilation were unchanged (Figure S7F-H). In the inflation challenge assay, we did not observe attenuation of the effect of inflation holding on either inspiratory or expiratory time (Figure 7H, S7I, and S7J). Thus, PSN7 neurons do not mediate the respiratory reflex induced by lung inflation. In summary, these results suggested that PSN7 neurons are also required for setting eupneic tidal volume, but unlike PSN6 neurons, they appear to do so by regulating inspiratory flow.

## Discussion

Combining organ-specific retrograde labeling and single-cell RNA-sequencing, here we provide a comprehensive, high-quality molecular cell atlas of vagal sensory neurons innervating the mouse lung. This atlas defines 10 molecular PSN subtypes, triple the number inferred from classical physiological studies, and likely comprising the full set. From our analysis of individual subtypes regarding their developmental lineages, myelination states, sensory and signaling gene profiles, terminal morphologies and locations, and response properties to mechanical challenges, we can integrate our atlas with prior studies of PSNs (Figure 8A). The whole transcriptomic profiles of individual subtypes uncover potential new functions of these neurons and the mediating signaling pathways, both in transmitting interoceptive information to the central nervous system and in directly interacting with cells in the lungs. Our initial characterization of the physiological functions of two *Piezo2*-expressing subtypes further reveal that breathing behavior is likely modulated by pulmonary sensory input in more diverse ways than previously appreciated.

**Figure 8.**
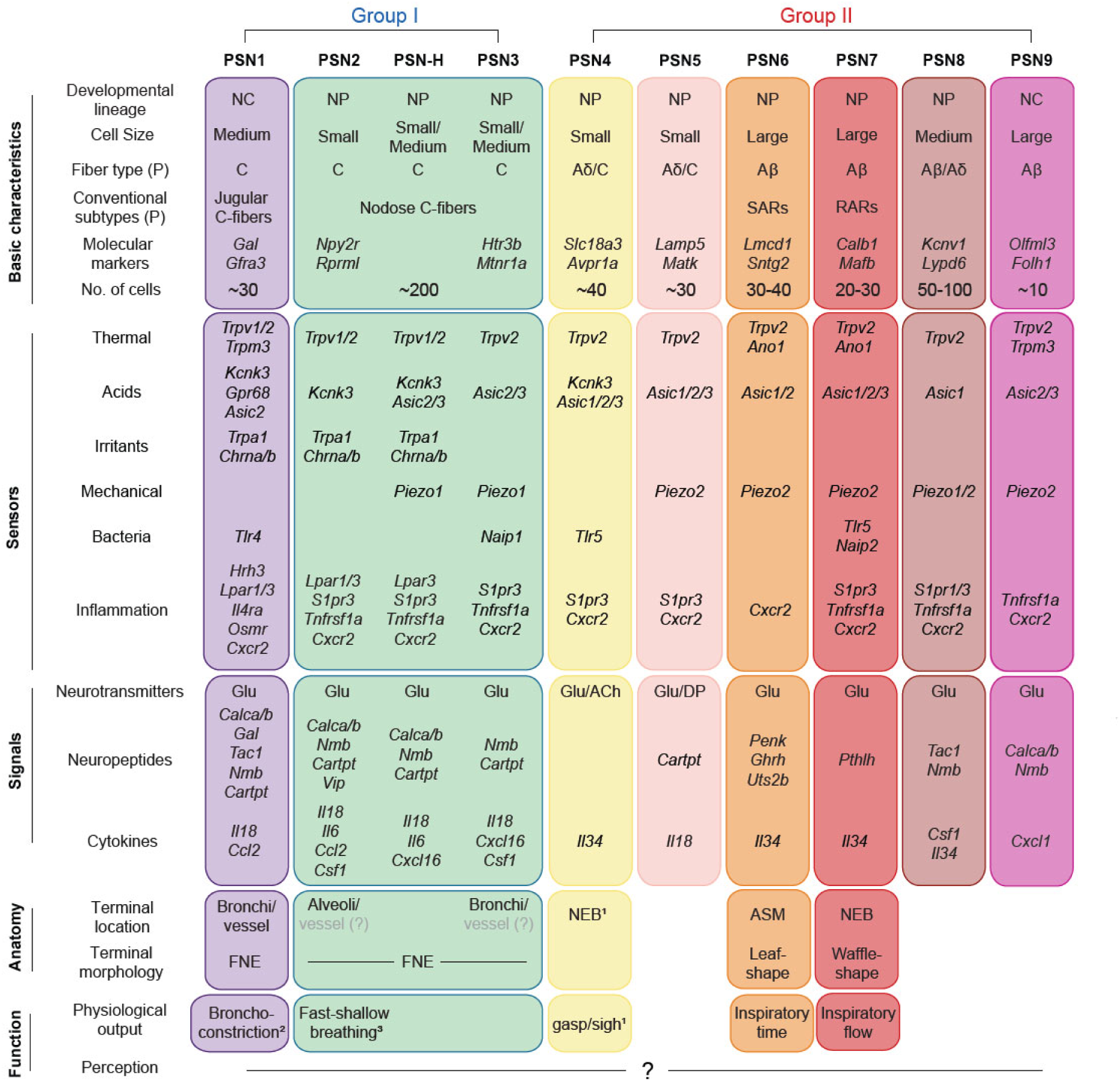
Summary of the molecular, anatomical, and functional diversity of vagal PSNs. Chart summarizing the molecular, anatomical, and functional diversity of PSN subtypes defined in this work integrated with prior studies. NC, neural crest lineage, NP, neural placode lineage, (P), predicted. Physiological outputs of PSN1, PSN2, and PSN4 are inferred by combining molecular identities recognized in this study and previous functional studies (1, Schappe et al. 2024; 2, Han et al. 2018; 3, Chang et al. 2015). Little is known about the perceptions generated by each PSN subtype.

We identified four subtypes, PSN1, 2, 3 and H, that represent the pulmonary C-fibers described by classical studies. PSN1 and PSN2 are two typical irritant-sensing C-fiber subtypes that express high levels of *Trpv1* and *Trpa1*. These two subtypes differ in developmental lineage (PSN1, neural crest; PSN2, neural placode) and receptive field (conducting airways vs. alveoli). They likely also differ in the physiological responses elicited by their activation. Activating PSN1 neurons with an agonist (Bam8-22) for PSN1-specific *Mrgprx1* (encode MrgprC11 in mouse), leads to reflex bronchoconstriction ^67,68^, whereas activating a subset of vagal sensory neurons expressing *Npy2r*, which includes PSN2 (and a subset of PSN-H), causes rapid and shallow breathing ^24^. Thus, signals activating receptors expressed by both subtypes, such as aldehydes in polluted air and inflammatory signal lysophosphatidic acid, would likely trigger distinct physiological responses in different lung compartments (conducting airways vs. alveoli). PSN3 appears to have similar terminal structures, locations, and distributions as PSN1, but unlike PSN1, it does not express irritant receptor *Trpa1* but expresses *Piezo1*, so is potentially mechanosensitive. Compared to PSN2, PSN3 neurons express lower levels of *Trpv1* transcripts and slightly higher levels of *Nefh*. Previous electrophysiological recording experiments had revealed a subpopulation of pulmonary C-fibers with slightly faster conduction velocity and insensitive to 1µM capsaicin, which fits the molecular profile of PSN3 ^69^. The other newly identified C-fiber subtype, PSN-H, is a curious molecular hybrid between PSN3 and PSN2. We did not identify any expressed genes unique to PSN-H, rather PSN-H neurons comprise a continuous molecular gradient between PSN2 and PSN3 identities, reminiscent of a few continuously heterogeneous neuronal populations observed in the brain ^70,71^. It will be important to determine the distribution and dynamics of PSN-H neurons *in vivo* to see if they form a spatial or functional gradient or represent dynamic intermediates between plastic PSN2 and PSN3 fates.

The other six PSN subtypes (Group II) include five putative mechanoreceptor subtypes (PSN5-9) expressing *Piezo2*. PSN6-9 are predicted to be myelinated and fast conducting A-fiber neurons, thus presumably include the classical SARs and RARs. We focused on PSN6 and PSN7 neurons and revealed their terminal structures, mechanical properties, and physiological functions in breathing regulation. PSN6 neurons target bronchial smooth muscle and respond to lung inflation, presumably by sensing smooth muscle stretch. They resemble classical SARs, and their function is to limit inspiratory time and provide homeostatic feedback on lung volume. PSN7 neurons, by contrast, terminate atop NEBs on the luminal (apical) side at bronchial branch points, and their function appears to be limiting inspiratory airflow. A small subset of these neurons exhibit the typical response property of RARs to lung inflation, suggesting that they are not lung volume sensors but rather report dynamic information. Given their anatomical features, they could potentially sense other types of mechanical forces generated in the airway lumen, such as mucus movement. The observation that not all neurons of either subtype respond to inflation challenge indicates that there is potentially further physiological heterogeneity among neurons of the same molecular subtype. One possibility is that neurons terminating at different locations in the lungs have different activation thresholds, or the property of their cells of contact, such as smooth muscle and neuroendocrine cells, determines if the neurons would respond or not. The different physiological outputs generated by PSN6 and PSN7 suggest that they synapse with separate populations of second-order neurons in the NTS, which in turn connect to different output circuits. One limitation of our functional interrogation experiments is that the ablations were not restricted to interoceptors of the lung, because neither *Calb1* nor *Mc4r* expression is restricted to WGA-labeled neurons after lung instillation. We directly assessed mechanical challenge-induced respiratory reflex, and the results supported our conclusion on the functional role of PSN6. For PSN7, static inflation appeared not to be an efficient way to activate these neurons, and their role on breathing regulation may not be reflected by diaphragm EMG recording. New strategies that target this population with both lung and subtype specificity are needed to confirm these conclusions.

The other Group II putative mechanoreceptor subtypes, PSN5, 8 and 9, may differ from PSN6 and PSN7 in both their sensory properties and physiological regulation functions. For instance, respiratory suppression at high levels of lung inflation was still observed after PSN6 ablation, suggesting other mechanoreceptor subtype(s) with a higher mechanical threshold mediate this effect, likely playing a role during vigorous exercise when tidal volume is much bigger. Also, PSN6 loss only affects inspiratory time but not expiratory time, suggesting other subtype(s) regulate the later breathing parameter. In addition to the surprising plethora and diversity of mechanosensory PSN subtypes, we also discovered a unique and novel subtype, PSN4, and its molecular features suggested a function related to hypoxia sensing. A recent work identified a similar molecular population that serves as the deflation receptor and sense airway collapse indirectly through pulmonary neuroendocrine cells ^39^. It would be interesting to explore if this subtype of neurons integrates multi-modality information in the lungs.

The sensory repertoire analysis revealed that each PSN subtype is capable of detecting multiple types of external and internal stimuli, including inflammatory signals and pathogens. This suggests that vastly different types of sensory information can be transmitted by the same PSNs. Some diverse stimuli may independently activate the same subtype and elicit the same physiological response, e.g., both hot air and nicotine induce bronchoconstriction ^41,45,72^. Other stimuli could be integrated, one potentiating or suppressing the PSNs’ response to another, e.g., prostaglandins heighten the sensitivity of pulmonary C-fibers to capsaicin ^73^ and hypercapnia suppresses the activity of pulmonary mechanoreceptors ^40^. Because subtypes can exhibit distinct firing patterns under different conditions, e.g., breathing-associated mechanical stimuli generate oscillatory firing whereas chemical stimuli cause continuous firing ^16^, there could also be sensory coding within a neuron if different firing patterns activate different cells and output pathways. Some of the inflammatory or pathogenic signals may not directly alter the electrical activity of PSNs, but rather induce transcriptional changes to alter their sensory repertoire or responsiveness ^68,74,75^.

A surprising finding from our analysis of the expression of secreted signals is that, in addition to immune-influencing neuropeptides, PSNs are also a source of many cytokines that can act on different immune cells and potentially form local signaling circuits. Some of these cytokines such as IL18 (asthma, COPD) and IL6 (COVID-19) are implicated in serious lung diseases (ref). Among PSNs, PSN1 stands out as a hub for bi-directional neuroimmune interactions with its high-level expression of multiple cytokines and cytokine receptors. PSN1 is a *Trpv1*^+^ subtype, and several studies have shown functions of *Trpv1*^+^ vagal sensory neurons in regulating immune responses in the lungs ^76–78^. Our transcriptomic atlas provides molecular candidates that mediate the interactions and regulatory functions. Moreover, we found that inflammatory receptor and cytokine expression is not restricted to *Trpv1*^+^ neurons. In fact, there are receptors and cytokines (*S1pr1*, *Il17rc*, *Csf1* and *Cxcl1*) selectively expressed by medium to large diameter putative mechanoreceptor subtypes (PSN8 and 9), suggesting their potentially distinct roles in neuroimmune interactions. Guided by predictions from the PSN expression profiles and initial characterization of the 10 subtypes, it is important to fully define experimentally what each PSN subtype senses, how these sensory inputs are integrated, and how that influences the local and central output signals and their impacts, in both health and disease.

Each internal organ has highly specific functions in orchestrating whole body physiology, and sensory neurons innervating individual organs are likely exposed to unique combinations of internal signals, as well as external stimuli if they innervate barrier organs, such as the lungs and the GI tract. Comparing the subtype compositions and molecular profiles of neurons innervating different internal organs would potentially reveal unique sensory populations or molecular mediators for organ-specific body-brain communications. A recent study made important progress on this topic, though not all internal organs were included and the completeness of coverage within each organ remains to be determined, highlighting the need for further efforts in the field ^27^. It seems likely that all organs are innervated by at least one subtype of danger/damage-sensing C-fibers plus varying numbers and diversity of subtypes serving in physiological feedback circuits as in the lungs but tailored during evolution to the function of each organ (e.g., nutrient and osmolarity sensors in the gut and the liver ^79–81^ and mechanoreceptors in the heart and vessels ^7,82^). Myelinated subtypes are likely reserved for functions in which rapid feedback with high temporal fidelity is critical ^83^, such as in breathing and cardiac regulation. Moreover, integrating the expression profiles of interoceptors and resident cells of other organs and comparing the interactomes of different organs should reveal both general interactions throughout the body and the ones specific for a particular organ. Such studies, ideally carried out in a variety of species including humans, will lay the groundwork for a new understanding of interoception and autonomic physiology, their evolution and their contributions to physical and mental health and disease.

## Supporting information

Key resources table

## Acknowledgements

We thank current and former members of the Krasnow lab for experimental support and comments on the manuscript, especially Joe Ouadah for help on scRNA-seq experiment, Kyle Travaglini for organizing and sharing scRNA-seq data of the mouse lung, Issac Domenech (Stanford Summer Research Program) and Andrea Yung for improving the smFISH protocol on vagal ganglion sections, Josh Head for assistant with whole-body plethysmography recordings, and Christin Kuo for discussions. We thank Zachary Knight, Nirao Shah and Eiman Azim for reagent sharing. We thank Ben Passarelli, Stanford Stem Cell Institute Genome Center, Stanford Functional Genomic Facility, and Shihong Gao from Janelia Scientific Computing for support on scRNA-seq experiments and following analysis. This work was supported by HHMI through both the investigator program and Janelia Research Campus, Damon Runyon postdoctoral fellowship (Y.L.), Stanford Child Health postdoctoral fellowship (Y.L.), and NHLBI (F31-HL137406) (A.J.D.). M.A.K. is a HHMI investigator.

## Author contributions

Y.L. performed all experiments. A.J.D. contributed to viral production and RNAscope ISH experiments. L.K. contributed to *in vivo* calcium imaging. Y.L. and M.A.K conceived and designed the experiments, interpreted data, and wrote the manuscript.

---

**Figure S1.**
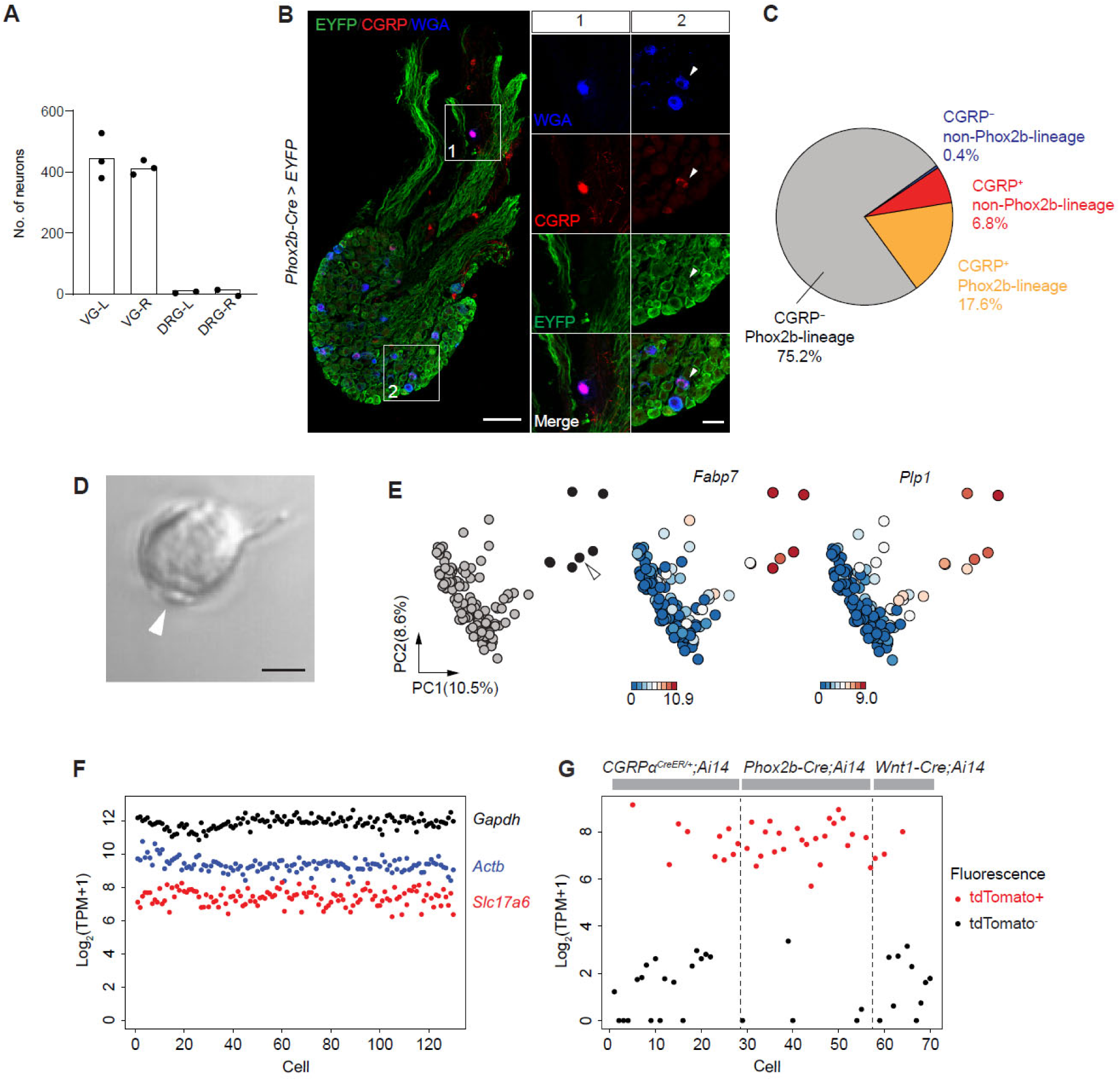
Single-cell RNA-seq of vagal PSNs, related to Figure 1. (A) Quantification of WGA-labeled neurons in vagal (VG) and dorsal root ganglia (DRG, C7-T6 collectively) on the side of the body indicated (L, left, R, right). Dots, values from individual mice; bars, mean values. (B) A vagal ganglion section from a *Phox2b-Cre;*Ai32/+ mouse with WGA labeling from the lungs, immunostained for indicated markers. Close-ups of boxed areas are shown in split panels on the right. Box 1, a CGRP^+^ Phox2b-lineage negative (EYFP^-^) WGA-labeled neuron. Box 2, a CGRP^+^ Phox2b-lineage positive (EYFP^+^) WGA-labeled neuron (arrowhead). Scale bars: 100 µm, 20 µm (insets). (C) Fractions of WGA-labeled vagal sensory neurons with indicated combinations of CGRP and Phox2b-lineage positivity (n=1122 scored neurons in 3 ganglia from 3 mice). (D) Brightfield image of a PSN with satellite glial cell(s) still attached (arrowhead) after ganglion cell dissociation. Scale bar: 10 µm. (E) PC plots showing 7 cells (black) separated from others (gray) when satellite glial cell-enriched genes were used for analysis. These cells expressed high levels of glial cell marker genes *Fabp7* and *Plp1*. Expression scale: log_2_(TPM+1) for range indicated. Arrowhead, cell shown in (D). (G) Sequencing results of housekeeping genes (*Gapdh*, *Actb*) and a sensory neuron marker gene (*Slc17a6*) across all cells passed quality control. High consistency and no drop-out indicate high quality expression profiles. (H) Sequencing results of *tdTomato* expression in cells picked from mice with genetic labeling as indicated. Red and black dots, cells with (red) and without (black) tdTomato fluorescence observed during cell picking. Note sequencing results were consistent with the fluorescence records. Low levels of tdTomato reads detected in some of the fluorescence-negative cells are likely due to low-level leaky expression of the reporter gene or contaminating mRNA released from dying or dead tdTomato-expressing cells, given that it was not observed in neurons from wild type mice.

**Figure S2.**
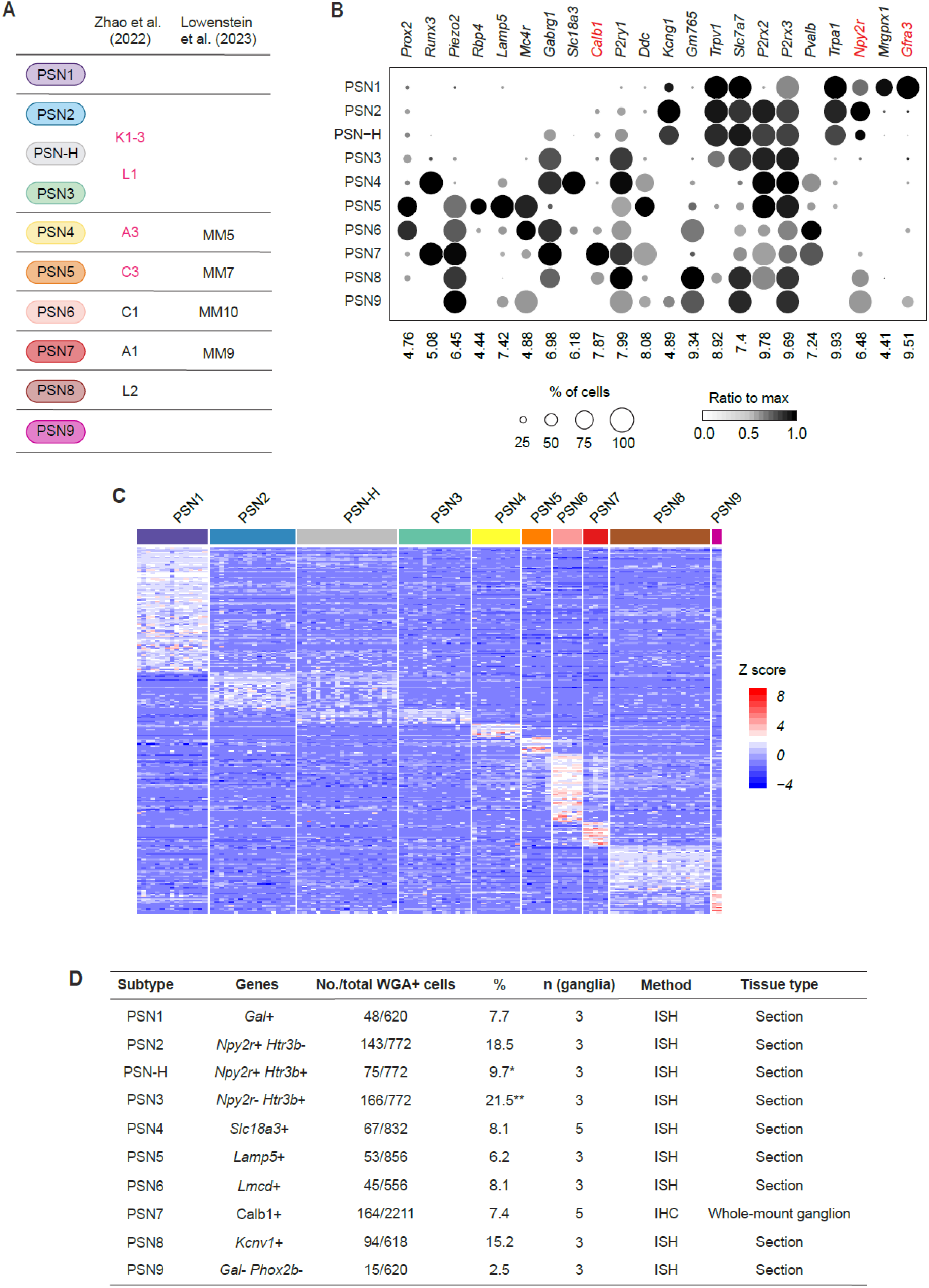
Molecular markers and abundances of PSN subtypes, related to Figure 2. (A) Matching PSN subtypes with vagal sensory clusters identified in two previously published scRNA-seq studies based on molecular marker expression shown in (B). Magenta clusters from Zhao et al. (2022) were identified as lung-innervating clusters. (B) Expression levels of genes used for matching PSN subtypes in this study with previously described vagal sensory clusters or subpopulations of vagal PSNs. Red highlights genes previously identified as markers of subpopulations of vagal PSNs and enriched in single PSN subtypes in this study. (C) Heatmap showing relative expression levels of 260 subtype-enriched genes across the 10 molecular subtypes. Z scores were calculated for individual genes across all cells. (D) Quantification of ISH and immunostaining results using probes/antibodies for the subtype selective genes. Calb1 was detected by immunostaining on whole-mount ganglia, therefore yielded a significantly higher number of total WGA^+^ cells. *, likely lower than the true fraction of PSN-H neurons since *Npy2r* is expressed in a subset of these neurons. **, likely higher than the true fraction of PSN3 neurons for the same reason.

**Figure S3.**
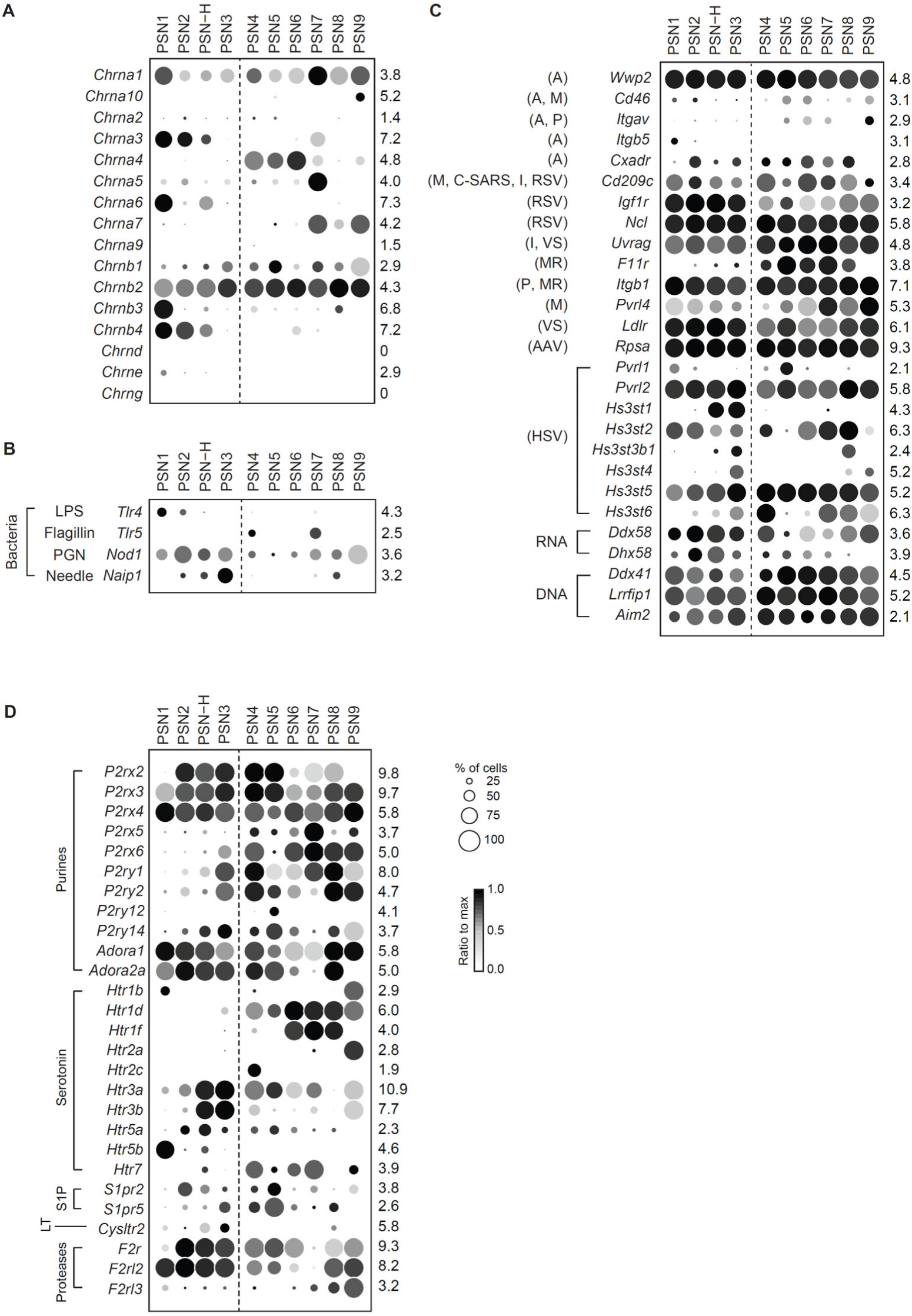
Functional receptors expressed in PSN subtypes, related to Figure 3. (A) Expression patterns of nicotinic acetylcholine receptor subunits across PSN subtypes. (B) Expression of pathogen pattern recognition receptors for bacterial cellular components. LPS, lipopolysaccharide; PGN, peptidoglycan. (C) Expression of entry receptors for respiratory viruses and intracellular nucleic acid sensors (magenta) across PSN subtypes. A, adenovirus; M, measles; P, parechovirus; C-SARS, SARS-coronavirus; I, influenza; RSV, respiratory syncytial virus; VS, vesicular stomatitis virus; MR, mammalian orthoreovirus; AAV, adeno-associated virus; HSV, herpes simplex virus. (D) Expression of additional receptors for inflammatory mediators. S1P, sphingosine-1-phosphate; PG, prostaglandins; LT, leukotrienes.

**Figure S4.**
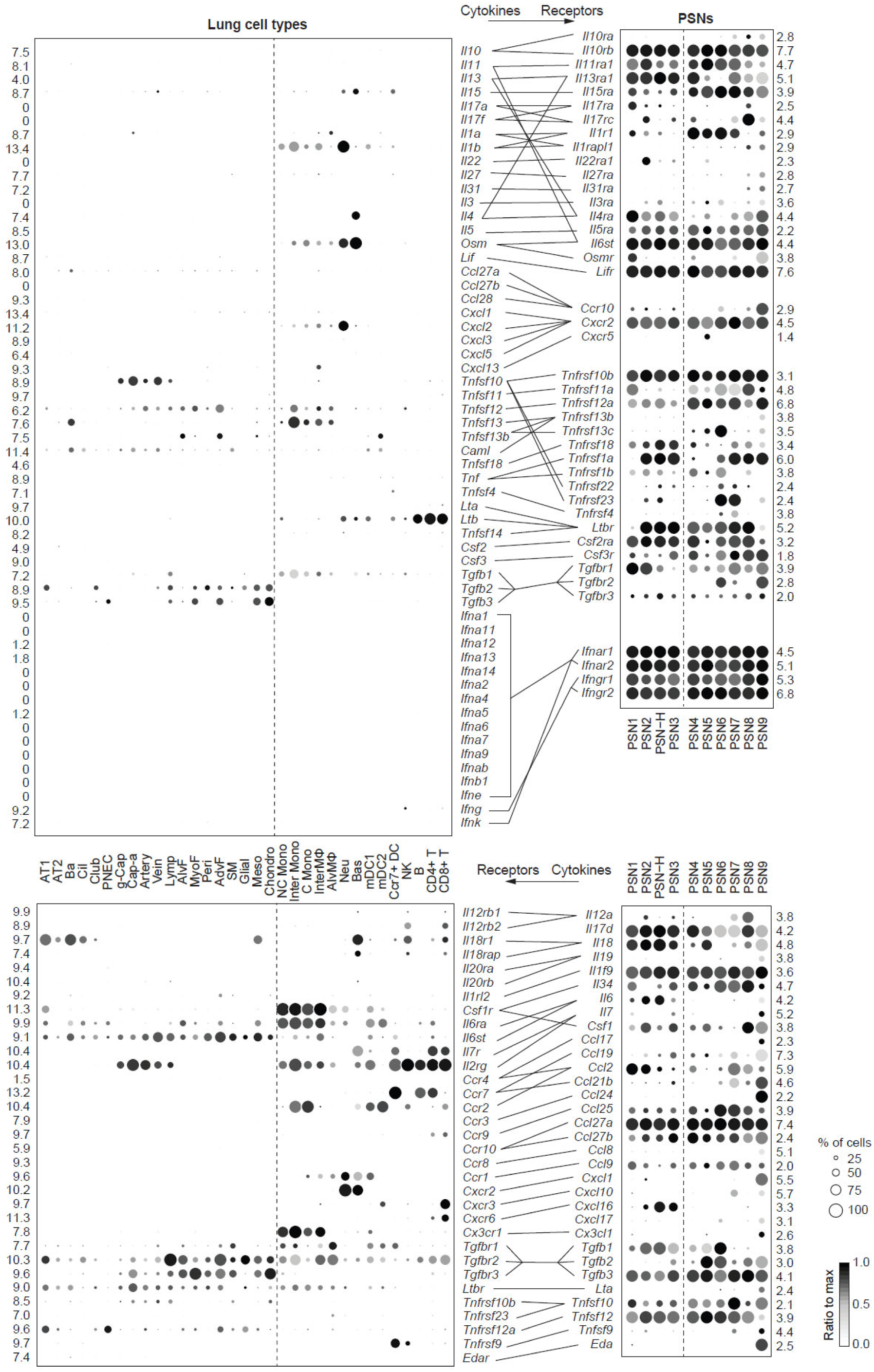
Local interactions between lung cells and PSNs, related to Figure 3. Predicted bi-directional cytokine signaling interactions between lung cells and vagal PSNs, inferred from expression of cytokine genes and the cognate receptor genes (connected by lines, arrows at top show the signaling direction). Full list of ligands and receptors screened is in Table S5, and genes expressed in >50% of neurons and a mean log2(TPM+1) > 1 in at least one PSN subtype are shown here. Dashed lines divide lung immune (right side) from non-immune (left) cell types (left dot plots), and Group I (left) from Group II (right) PSN subtypes (right dot plots). AT1, alveolar type 1 cells; AT2, alveolar type 2 cells; Ba, basal cells; Cil, ciliated cells; Club, club cells; PNEC, pulmonary neuroendocrine cells; Cap, general capillary endothelial cells (g-cap); Cap-a, capillary aerocytes; Artery, arterial endothelial cells; Vein, venous endothelial cells; Lymp, lymphatic endothelial cells; AlvF, alveolar fibroblasts; MyoF, myofibroblasts; Peri, pericytes; AdvF, adventitial fibroblasts; SM, smooth muscle cells; Glial, peripheral glial cells; Meso, mesothelial cells; Chondro, Chondrocytes; mDC1, myeloid dendritic type 1 cells; mDC2, myeloid dendritic type 2 cells; Ccr7+ DC, Ccr7+ dendritic cells; NC Mono, non-classical monocytes; inter Mono, intermediate monocytes; C Mono, classical monocytes; B, B cells; CD4+ T, CD4+ T cells; CD8+ T, CD8+ T cells; NK, natural killer cells; Neu, neutrophils; Bas, Basophils; InterMɸ, interstitial macrophages; AlvMɸ, alveolar macrophages.

**Figure S5.**
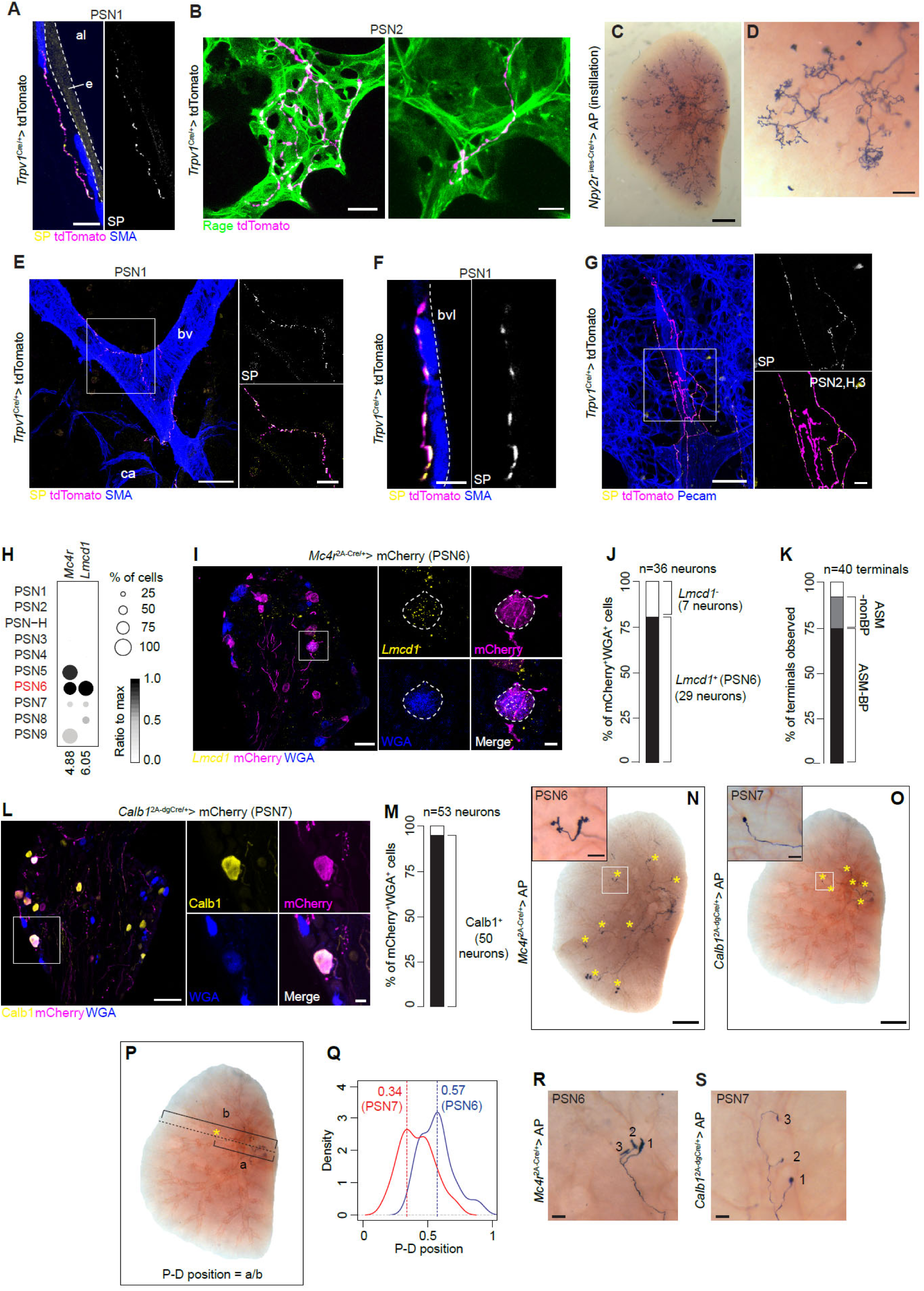
Terminal locations and morphologies of PSN subtypes, related to Figure 4. (A) Immunostaining of a lung section from *Trpv1*^Cre/+^ mice injected with AAV-CAG-DIO-tdTomato into the vagal ganglia, showing a SP+tdTomato+ fiber (PSN1) runs beneath the airway smooth muscle stained for smooth muscle actin (SMA). Note gaps between smooth muscle fibers on conducting airways, which likely allow sensing of environmental chemicals just penetrating epithelium or molecules secreted by epithelial cells. e, epithelium (outlined); al, airway lumen. Scale bar: 20 μm. (B) Examples of alveolar terminals (likely PSN2) elaborating on the surface of an air sac (left) and between air sacs (right, more common). Scale bars: 10 μm. (C, D) Whole-mount alkaline phosphatase (AP) staining of lung lobes from *Npy2*^rires-Cre/+^ mice intratracheally instilled with the Cre-dependent AP virus (labels PSN2). Note in D, all branches of the labeled fibers terminate in the alveolar region but not on the conducting airways. Scale bars: 1 mm (C), 0.1 mm (D). (E, F) SP+tdTomato+ fibers (PSN1) terminate on pulmonary blood vessels (bv). ca, conducting airway. F shows a fiber running beneath the vascular smooth muscle. Scale bars: 100 μm (E), 20 μm (E inset), 5 μm (F). (G) SP-tdTomato+ fibers (PSN2, H, or 3) also terminate on pulmonary blood vessels (endothelial cells stained by Pecam, blue). Scale bars: 50μm, 10 μm (inset). (H) Expression patterns of *Mc4r* and PSN6 marker *Lmcd1* across PSN subtypes. (I) ISH of PSN6 subtype marker *Lmcd1* with mCherry and WGA double immunostaining on a vagal ganglion section from a *Mc4r*^2A-Cre/+^ mouse injected with AAV-Syn-DIO-hM3Dq-mCherry into the ganglion. Close-up of the boxed area is shown in right panels with channels split; outlined neuron is an example of labeled PSN6 neuron. Scale bars: 50μm, 10μm (inset). (J) Quantification of PSN6 labeling specificity (serial sections of two ganglia from two *Mc4r*^2A-Cre/+^ mice). Lmcd1-labeled neurons are likely PSN5 neurons. (K) Locations of labeled termini in *Mc4r*^2A-Cre/+^ mice injected with reporter virus into the ganglia. Quantifications were done by examining all vibratome sections of 4 entire left lobes from 4 mice. ASM, airway smooth muscle; BP, branch point. (L) Immunostaining of PSN7 marker Calb1 with mCherry and WGA on a vagal ganglion section from a *Calb1*^2A-dgCre/+^ mouse injected with AAV-Syn-DIO-hM3Dq-mCherry into the ganglion. Close-up of the boxed area with a labeled PSN7 neuron is shown in right panels with channels split. Scale bars: 50 μm, 10 μm (inset). (M) Quantification of PSN7 labeling specificity (serial sections of 3 ganglia from 2 *Calb1*^2A-dgCre/+^ mice). (N, O) Whole-mount AP staining of right cranial lung lobes from *Mc4r*^2A-Cre/+^ (left) and *Calb1*^2A-dgCre/+^ (right) mice injected with AAV-CMV-FLEX-PLAP into the vagal ganglia. Asterisks mark termini in each lobe. Insets are close-ups of boxed regions showing terminal morphology. Scale bars: 1 mm, 0.1 mm (insets). (P) Method of calculating proximal-distal (P-D) position of PSN termini. A line between the primary bronchus entry point into the lobe and a given terminal was drawn and extended to the edge of the lobe. Terminal P-D position is the ratio of the segment length between the entry point and the terminal to the total length of the line. (Q) Kernel density plot showing distributions of PSN6 (blue) and PSN7 (red) terminal P-D positions (n=29 PSN6 termini, n= 36 PSN7 termini). Note PSN7 termini are distributed more proximally than PSN6 termini. (R, S) Close-ups from whole-mount AP staining showing termini of single PSN6 (S) and PSN7 (T) fibers. Individual termini are numbered. Scale bar: 0.1 mm. We cannot exclude the possibility that these axons also terminate in other lung lobes or organs.

**Figure S6.**
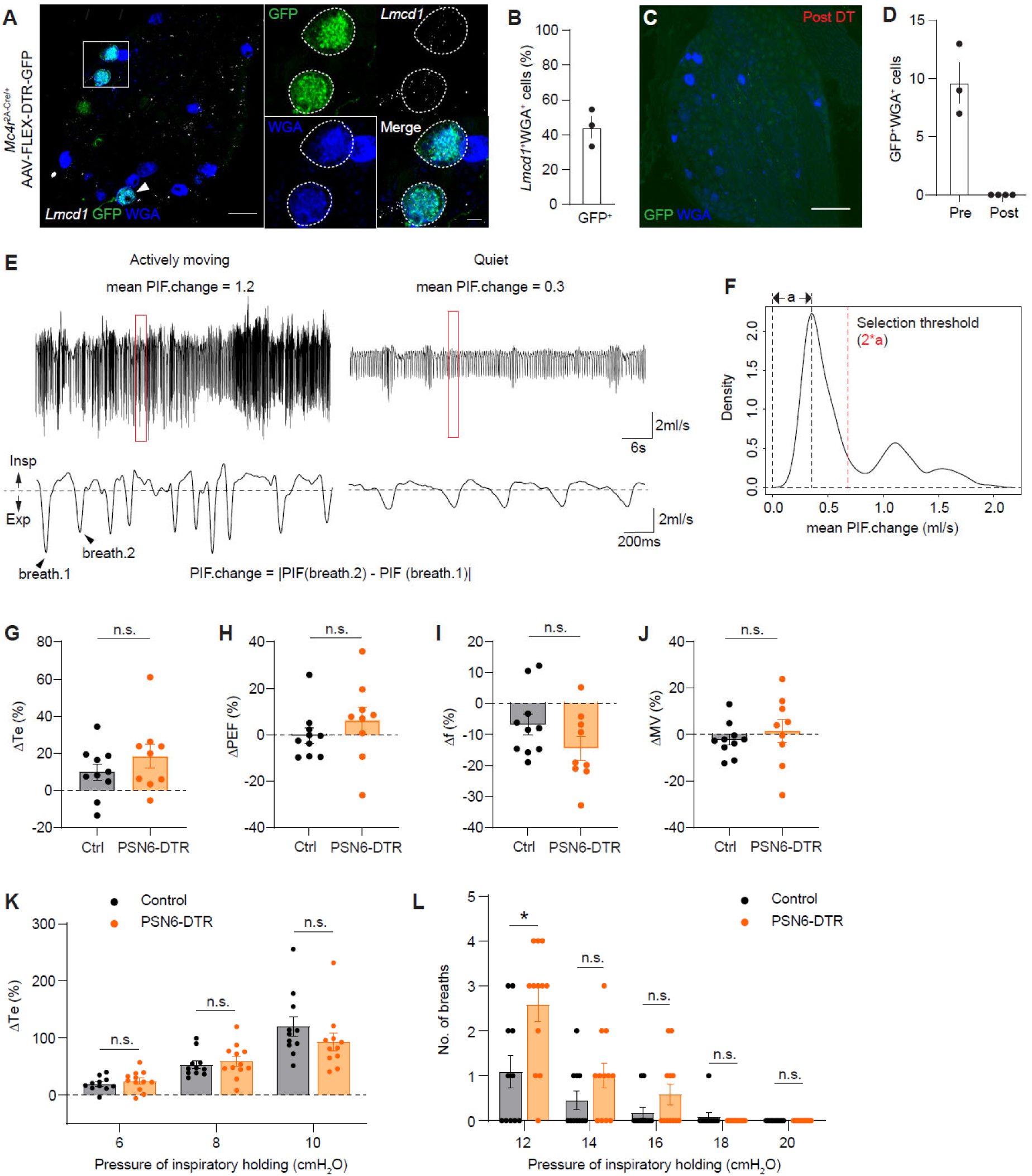
PSN6 ablation and breathing changes, related to Figure 6. (A) ISH of PSN6 subtype marker *Lmcd1* with GFP and WGA double immunostaining on a thin vagal ganglion section from a *Mc4r*^2A-Cre/+^ mouse injected with AAV-CBA-FLEX-DTR-GFP into the ganglia. WGA was instilled into the lung to label PSNs. Panels at right are split channels of the boxed region showing two DTR-GFP expressing PSN6 neurons. Arrowhead, another DTR-GFP expressing PSN6 neuron. Scale bars: 50 μm and 10 μm (inset). (B) Quantification of the percentage of *Lmcd1*+WGA+ PSN6 neurons expressing DTR-GFP in PSN6-DTR mice (n=57 *Lmcd1*+WGA+neurons scored on 3 ganglia from 3 mice, with each dot representing the result from all serial sections of one ganglion). (C) Immunostaining for GFP and WGA in a ganglion section from a PSN6-DTR mouse three days after DT injection. (D) Quantification of DTR-GFP expressing neurons in vagal ganglia of PSN6-DTR mice before and after DT injection (each dot represents the result from all serial sections of one ganglion). (E) Selection of quiet breathing periods. Representative airflow traces during actively moving (left) or quiet (right) phases, recorded by whole body plethysmography. Portions of traces highlighted by red boxes are enlarged below. PIF.change is the absolute difference of peak inspiratory flows of two adjacent breaths. Mean PIF.change is the average PIF.change in one minute period. Note the large difference in this parameter between active (1.2 ml/s) and quiet (0.3 ml/s) breathing phases. Dashed lines, zero flow; Insp, inspiration; Exp, expiration. (F) Kernel density plot showing distribution of mean PIF.change for every minute over the entire recording period. Time periods with mean PIF.change less than twice the PIF.change at the first peak (a) were selected as quiet breathing periods. (G-J) Changes in expiratory time (Te, panel G), peak expiratory flow (PEF, panel H), respiratory frequency (f, panel I), and minute ventilation (MV, panel J) distribution modes after DT injection (post), normalized to before DT injection (pre) values, in control and PSN6-DTR mice. (K) Changes in Te in control and PSN6-DTR mice after DT injections. ΔTe represents the change in the average time between EMG bursts during the first 2 seconds of inspiratory holdings at a given pressure, normalized to the average time between EMG bursts during the first 2 seconds of expiratory holding. Two-way ANOVA with sidak multiple comparison correction. (L) Number of breaths during the first 2s of inspiratory holdings at different pressures in control and PSN6-DTR mice after DT injections. *p=0.0496, two-way ANOVA with sidak multiple comparison correction.

**Figure S7.**
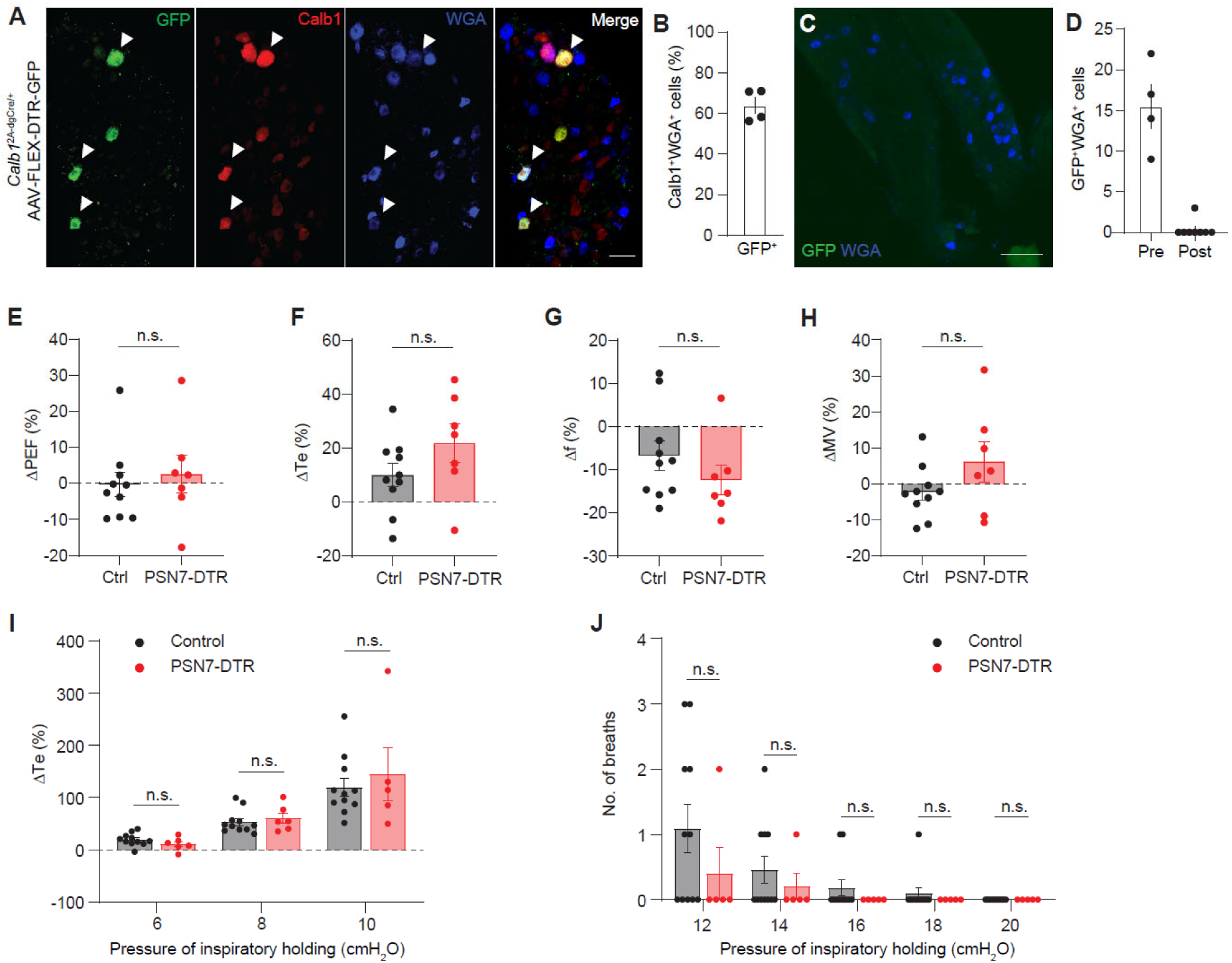
PSN7 ablation and breathing changes, related to Figure 7. (A) Immunostaining of PSN7 subtype marker Calb1 along with GFP and WGA on a thin vagal ganglion section from a *Calb1*^2A-dgCre/+^ mouse injected with AAV-CGA-FLEX-DTR-GFP into the ganglion and treated with TMP. WGA was instilled into the lung to label PSNs. Arrowheads, DTR-GFP expressing PSN7 neurons. Scale bar: 50 µm. (B) Quantification of the percentage of Calb1^+^WGA^+^ PSN7 neurons expressing DTR-GFP in PSN7-DTR mice (n= 95 Calb1^+^WGA^+^ neurons scored from 4 ganglia of 2 mice, with each dot representing result from all serial sections of one ganglion). (C) Immunostaining for GFP and WGA on a ganglion section from a PSN7-DTR mouse three days after DT injection. (D) Quantification of DTR-GFP expressing PSNs in vagal ganglia of PSN7-DTR mice before (pre) and after (post) DT injection (each dot represents the result from all serial sections of one ganglion). (E-H) Changes in expiratory time (Te, panel G), peak expiratory flow (PEF, panel H), respiratory frequency (f, panel I), and minute ventilation (MV, panel J) distribution modes after DT injection (post), normalized to before DT injection (pre) values, in control and PSN7-DTR mice. (I) Changes in Te in control and PSN7-DTR mice after DT injections. ΔTe represents the change in the average time between EMG bursts during the first 2 seconds of inspiratory holdings at a given pressure, normalized to the average time between EMG bursts during the first 2 seconds of expiratory holding. Two-way ANOVA with sidak multiple comparison correction. (J) Number of breaths during the first 2s of inspiratory holdings at different pressures in control and PSN7-DTR mice after DT injections. Two-way ANOVA with sidak multiple comparison correction.

