## Supplementary material for "Molecular, anatomical, and functional organization of lung interoceptors": Key resources table

| **REAGENT or RESOURCE** | **SOURCE** | **IDENTIFIER** |
| --- | --- | --- |
| **Antibodies** | | |
| Chicken anti-GFP (1:500) | Abcam | Cat# 13970 |
| Rabbit anti-Calb1 (1:10,000) | Swant | Cat# CB-38 |
| Goat anti-mCherry (1:1000) | Scigen | Cat# AB0040-200 |
| Rabbit anti-RFP (1:1000) | Rockland | Cat# 600-401-379 |
| Rabbit anti-WGA (1:1000) | Sigma-Aldrich | Cat# T4144 |
| Goat anti-WGA (1:500) | Vector Lab | Cat# AS-2024 |
| Rat anti-substance P (1:500) | Millipore | Cat# MAB356 |
| Rabbit anti-CGRP (1:1000) | EuroProxima | Cat# 2263BGP470-1 |
| Mouse anti-SMA Alexa 488 conjugated (1:200) | Sigma | Cat# F3777 |
| Hamster anti-Pecam (1:500) | Bio-rad | Cat# MCA1370Z |
| Rat anti-Rage (1:500) | R&D | Cat# MAB1179 |
| **Bacterial and Virus Strains** | | |
| AAV9-hSyn-DIO-hM3Dq-mCherry | Gift from Dr. Zachary Knight | N/A |
| AAV1-CMV-FLEX-PLAP | Gift from Dr. Nirao Shah | N/A |
| AAVrg-CMV-FLEX-PLAP | Janelia Research Campus | N/A |
| AAV9-CBA-DIO-DTR-GFP | Janelia Research Campus | N/A |
| AAV9-CAG-FLEX-tdTomato | Penn Vector Core/Addgene | Cat# 51503 |
| **Chemicals, Peptides, and Recombinant Proteins** | | |
| Diphtheria toxin | List Labs | Cat# 150 |
| Trimethoprim | Sigma-Aldrich | Cat# 7883 |
| Wheat Germ Agglutinin (no conjugation) | Vector Lab | Cat# L-1020 |
| Wheat Germ Agglutinin (conjugated with Alexa fluor 488, Alexa fluor 555, Alexa fluor 594, Alexa fluor 647) | Thermo Fisher | Cat# W11261, W32464, W11262, W32466 |
| Wheat Germ Agglutinin (conjugated with CF®594) | Biotium | Cat# 29023 |
| **Critical Commercial Assays** | | |
| SMART-Seq v3 for mRNA-seq | Takara | Cat# 634849 |
| Nextera XT DNA Library Preparation Kit (96 samples) | Illumina | Cat# FC-131-1096 |
| Nextera XT Index Kit (96 indexes, 384 samples) | Illumina | Cat# FC-131-1002 |
| RNAscope Fluorescent Multiplex Reagent Kit | Advanced Cell Diagnostics | Cat# 320850 |
| **Deposited Data** | | |
| scRNA-seq data of adult mouse vagal pulmonary sensory neurons | This paper | GSE186180 |
| **Experimental Models: Organisms/Strains** | | |
| Mouse: Wild type: C57BL/6J | The Jackson Laboratory | JAX: 00064 |
| Mouse: Wnt1-Cre: H2az2^Tg(Wnt1-cre)11Rth^ | The Jackson Laboratory | JAX: 003829 |
| Mouse: Phox2b-Cre: B6(Cg)-Tg(Phox2b-cre)3Jke | The Jackson Laboratory | JAX: 016223 |
| Mouse: CGRP^CreER^: Calca^tm1.1(cre/ERT2)Ptch^ | Pao-Tien Chuang, UC San Francisco | MGI: J:190366 |
| Mouse: Mc4r-2A-Cre: Mc4r^tm3.1(cre)Lowl^ | The Jackson Laboratory | JAX: 030759 |
| Mouse: Calb1-2A-dgCre: B6.Cg-Calb1^tm1.1(folA/cre)Hze^ | The Jackson Laboratory | JAX: 023531 |
| Mouse: Trpv1-Cre: B6.129-Trpv1^tm1(cre)Bbm^ | The Jackson Laboratory | JAX: 017769 |
| Mouse: Ai14: B6.Cg-Gt(ROSA)26Sor^tm14(CAG-tdTomato)Hze^ | The Jackson Laboratory | JAX: 009714 |
| Mouse: Ai32: B6.Cg-Gt(ROSA)26Sor^tm32(CAG-COP4*H134R/EYFP)Hze^ | The Jackson Laboratory | JAX: 024109 |
| Mouse: Npy2r^ires-Cre^: Npy2r^tm1.1(cre)Lbrl^ | The Jackson Laboratory | JAX: 029285 |
| Mouse: Ai95D: B6J.Cg-Gt(ROSA)^26Sortm95.1(CAG-GCaMP6f)Hze^/MwarJ | The Jackson Laboratory | JAX: 028865 |
| Mouse: Ai148: B6.Cg-^Igs7tm148.1(tetO-GCaMP6f,CAG-tTA2)Hze^/J | The Jackson Laboratory | JAX: 030328 |
| Mouse: Ai65D: B6;129S-Gt(ROSA)^26Sortm65.1(CAG-tdTomato)Hze^/J | The Jackson Laboratory | JAX: 021875 |
| **Oligonucleotides** | | |
| RNAscope probe to mouse *Gal* | Advanced Cell Diagnostics | Cat#: 400961-C2 |
| RNAscope probe to mouse *Npy2r* | Advanced Cell Diagnostics | Cat#: 315951 |
| RNAscope probe to mouse *Htr3b* | Advanced Cell Diagnostics | Cat#: 497541-C3 |
| RNAscope probe to mouse *Slc18a3* | Advanced Cell Diagnostics | Cat#: 448771-C3 |
| RNAscope probe to mouse *Lamp5* | Advanced Cell Diagnostics | Cat#: 451071-C2 |
| RNAscope probe to mouse *Lmcd1* | Advanced Cell Diagnostics | Cat#: 484761 |
| RNAscope probe to mouse *Kcnv1* | Advanced Cell Diagnostics | Cat#: 508291-C3 |
| RNAscope probe to mouse *Phox2b* | Advanced Cell Diagnostics | Cat#: 407861-C3 |
| RNAscope probe to mouse *Piezo2* | Advanced Cell Diagnostics | Cat#: 439971 |
| **Recombinant DNA** | | |
| pAAV-CBA-DIO-DTR-GFP | Gift from Dr. Eiman Azim (Azim et al., 2014) | N/A |
| pAAV-CMV-FLEX-PLAP | Addgene | Cat#: 80422 |
| **Software and Algorithms** | | |
| kallisto v0.43.0 | Lior Pachter | RRID:SCR_016582 |
| Scde v1.2.1 | Peter Kharchenko (Kharchenko et al., 2014) |  |
| Jackstraw v1.3 | Neo chung |  |
| SigClust v1.1.0 | Hanwen Huang |  |
| Seurat v3.0 | Rahul Satija | RRID:SCR_007322 |
| R v3.5.3 | R Foundation for Statistical Computing | RRID:SCR_001905 |
| R studio v1.1.463 | Rstudio | RRID:SCR_000432 |
| ImageJ | NIH | RRID:SCR_003070 |
| Prism 10 | Graphpad | RRID:SCR_002798 |
| Other | | |

Azim, E., Jiang, J., Alstermark, B., and Jessell, T.M. (2014). Skilled reaching relies on a V2a propriospinal internal copy circuit. Nature *508*, 357-363. 10.1038/nature13021.

Kharchenko, P.V., Silberstein, L., and Scadden, D.T. (2014). Bayesian approach to single-cell differential expression analysis. Nat. Methods *11*, 740-742. 10.1038/nmeth.2967.
