## Supplementary material for "Molecular, anatomical, and functional organization of lung interoceptors": Materials and Methods

**Resource availability**

Further information and requests for resources and reagents should be directed to Yin Liu and Mark A. Krasnow. All unique/stable reagents generated in this study are available from the corresponding authors upon proper request.

**Animals**

Mouse lines used in this study are listed in the Key Resources Table. Wild type (C57BL/6) mice were obtained from Jackson Laboratories. All studies were carried out on a mixture of adult (at least 6-week-old) male and female mice, with comparable numbers of each sex allocated to experimental groups. Mice were maintained in 12 hour light/dark cycle with food and water provided *ad libitum*. All animal husbandry, maintenance, and experiments were performed in accordance with Stanford University’s IACUC-approved protocols (APLAC 9780, 28836) and HHMI/Janelia Research Campus IACUC-approved protocol (22-0231).

**Retrograde labeling of vagal PSNs**

Mice were anesthetized with 2.5~3% isoflurane in induction chamber for at least 15 min and then placed on an intubation stand (Rodent Tilting Workstand, Hallowell EMC). Light from a microscope illuminator was placed gently against the neck. The mouse’s mandible was lifted, and tongue was extended with a Q-tip. Vocal cord was visualized with a speculum and an intubation tube (1 inch 20G disposable catheter, Fisher Scientific) was inserted into the trachea with a guide wire (Mouse Intubation Pack, Hallowell EMC). After removing the guide wire, 50 µl of 1mg/ml fluorescent conjugated WGA was instilled into the lung during natural breathing. For double labeling from the lung, each of the fluorescent conjugated WGAs (Alexa488 and Alexa555) was instilled separately one day apart. Two other commonly used tracers, DiIC_18_(3) (1mg/ml in 10% DMSO/90% saline, sonicated) and fluorescent conjugated CTB (1 mg/ml), were also tested, but they labeled fewer neurons than WGA (data not shown).

**Retrograde labeling of vagal stomach sensory neurons**

Mice were anesthetized with 1.5% isoflurane via nose cone (after induction with 2-3% isoflurane). A horizontal incision was made in abdominal skin, followed by an incision in the abdominal muscle to expose the stomach. Fluorescent conjugated WGA (5 ul of 5 mg/ml solution) was injected into the wall of the dorsal stomach (3~6 sites) using glass pipettes connected to an aspirator tube (Sigma, A5177).

**Single cell transcriptional profiling**

Mice used for collecting single PSNs for transcriptional profiling were adults 6-12 weeks old.

*Vagal ganglion dissociation*

Our protocol was modified from one previously described (Dubuis et al., 2013). Only one mouse was used for dissociation each day (two ganglia yield enough cells for picking, and the source animal for each cell was recorded so that transcriptomic variance introduced by individual mice could be analyzed and excluded during cluster analysis). Four to seven days after retrograde labeling, mice were euthanized with CO_2_ and transcardially perfused with cold Hank’s balanced salt solution (HBSS) (without Ca^2+^ and Mg^2+^). Vagal ganglia were quickly dissected after perfusion in cold buffered HBSS (10 mM HEPES, pH7.2-7.5) and each was cut into 2~3 pieces. Ganglion tissue was then digested with papain solution at 20 U/ml (Worthington LS003124, in buffered HBSS containing 0.5 mM EDTA, 1.5 mM CaCl2, 0.4 mg/ml L-cysteine, pre-warmed for 30 min for activation) for 10 min at 37°C, and then collagenase/dispase solution (collagenase IV, 300 U/ml, Worthington LS004186; dispase, 1mg/ml, Thermo Fisher Scientific 17105-041, in HBSS, pre-warmed) for 30 min at 37°C with tube inversion to agitate every 10 min. After centrifuging at 400 g for 4 min, tissue pellet was resuspended in 1 ml L15 medium (with 10mM HEPES pH 7.2-7.5 and 10% fetal bovine serum) and triturated using fire-polished Pasteur pipettes. A Percoll gradient spin was used to clear axonal debris and glial cells by layering the cell suspension onto a buffered L15 medium (10 mM HEPES)/Percoll (4:1) solution and then centrifuging for 9 min at 400 g. The cell pellet was resuspended in 2 ml buffered L15 medium, centrifuged again for 3 min at 750 g, and resuspended in 200 µl suspension medium (L15 medium, 10mM HEPES pH 7.2-7.5, 1mg/ml BSA, and 0.1% Pluronic F-68). We tested this protocol for satellite glial dissociation by placing the cells onto poly-d-lysine coated cover glasses, fixing with 4% PFA in PBS, and then staining for DAPI; less than 10% of single neurons were associated with additional nuclei on their side, indicating nearly complete removal of glial cells. The protocol is optimized for the nodose side of the vagal ganglion, and the jugular side (neural crest derived) was likely over-digested; only ~3% of WGA labeled cells were neural crest lineage after dissociation (assessed as tdTomato^+^ from *Wnt1-Cre*; *Ai14/+*mice or tdTomato^-^ from *Phox2b-Cre*; *Ai14/+* mice).

*Single cell picking*

The single cell suspension was placed on ice, and cell picking was carried out within 2 hours after completion of ganglion dissociation to maximize cell health. Drops (2 µl) of cell suspension were placed on a petri dish and examined with an inverted fluorescent microscope to identify WGA-488 labeled sensory neurons. Individual labeled neurons were collected using a custom glass micropipette with inside filament (WPI TW100F-4) connected to an aspirator tube (Sigma A5177). The micropipette was made with a pipette puller (Sutter instrument, P-87) and the tip fabricated by the “glass-on-glass” method (Pipette cookbook) to produce a 50-100 micron diameter opening. Liquid (0.3 µl or less) containing a single labeled cell was aspirated into the pipette by capillary action and then transferred to a fresh, ice-cold 2 µl drop of suspension medium. The cell was picked again with a fresh glass micropipette, transferred to a 0.2 ml PCR tube containing 4 µl of lysis buffer, and then immediately frozen on dry ice. Transfer to fresh medium and the second round of picking was performed to ensure purity. A micrograph was taken of each cell prior to the second pick, and later used to measure cell size. Single cell lysates were stored at -80°C before proceeding to cDNA synthesis and amplification.

*cDNA synthesis and amplification*

cDNA synthesis and amplification of single vagal PSNs were carried out using the SMART-Seq v3 Ultra Low Input RNA Kit (Clontech). To minimize amplification bias, cDNA amplification was limited to 14-15 cycles. The concentration and quality of cDNA was examined by Bioanalyzer using the High Sensitivity DNA kit (Agilent 5067-4626), and any sample with <30 pg/µl cDNA was discarded (1 of 137).

*Library preparation, sequencing, and raw data processing*

Our method for constructing sequencing libraries has been described (Ouadah et al., 2019). Briefly, amplified cDNA was diluted to 30~150pg/µl range and followed by tagmentation and amplification with index primers using the Nextera XT DNA Library Preparation Kit (Illumina FC-131-1096) and Index Adapters (Illumina, FC-131-1002). These steps followed the C1 system for mRNA Seq Protocol (Fluidigm). Library concentration and quality was examined by Bioanalyzer using the High Sensitivity DNA kit (Agilent 5067-4626), and sequencing was carried out on NextSeq 500 (Illumina) with 75 bp paired-end reads. Raw reads were aligned to the NCBI RefSeq mouse transcriptome (modified to exclude model XR_/XM_ sequences and include the tdTomato sequence) using *kallisto* with default settings (Bray et al., 2016). Read count and TPM values from splice isoforms of the same genes were collapsed, and for quality control cells with less than 2,000,000 paired-end reads that mapped to endogenous genes or less than 10,000 endogenous genes with mapped paired-end reads were excluded from the following analysis. Both raw reads and TPM counts were deposited to Gene Expression Omnibus (GSE186180).

**scRNA-seq data analysis**

*Identification and exclusion of glial-contaminated cells*

The TPM table was log-transformed (log_2_(TPM+1)) and scaled, and PCA was performed using 305 glial cell-enriched genes (Li et al., 2016). Cells were plotted along the first two PCs and 7 cells clearly separated from the rest (Figure S1F), which were considered glial-contaminated and eliminated from the following cluster analysis.

*Iterative cluster analysis*

Our algorithm is modified from one previously described (Tasic et al., 2016). For each round of iterative clustering, the following steps were performed:

1. Mean expression and variance were calculated based on log_2_(TPM+1) values for all endogenous genes.

2. Highly variable genes with different combinations of mean expression (1 to 3 with 0.5 increment) and variance (1 to 3 with 0.5 increment) thresholds were selected (25 sets). Step 3 to step 5 were carried out for each set.

3. PCA was performed using highly variable genes (*prcomp* function) and significant PCs were selected using the *Jackstraw* package. When the number of highly variable genes was smaller than the number of cells, data was randomized for each column (cell) and PCA repeated 100 times. Eigen values of each PC from the original PCA were compared with eigen values of PC1 from the permuted PCAs using a t-test, and PCs with multiple comparison adjusted p-value <0.05 were selected as significant PCs.

4. Euclidian distances were calculated using the cell loadings from the significant PCs and followed by hierarchical clustering using the “ward. D2” method.

5. Only the first split from hierarchical clustering was taken into consideration and the significance level of the split was calculated using *SigClust* package.

6. Among the 25 sets of highly variable genes, if no p-value from step 5 was <0.05, clustering was terminated.

7. If one or more p-values from step 5 was <0.05, the set of highly variable genes that resulted in the lowest p-value was selected and randomly subsampled (80%), and steps 3 to step 5 repeated 100 times. Cells that clustered into the same group in >90% of the events were accepted as a stable cluster, whereas cells that switched between groups in >10% of the events were labeled as outliers. When a group of cells (≥5 cells) switched their group identity together in >90% of the events, they were accepted as a third cluster.

8. Clusters comprising less than 5 cells or cells that all came from one animal were not promoted to the next round or accepted as final clusters.

9. After the final clusters were obtained, each cluster was cross-validated with each of the other clusters (except those that were resolved together at the final round of iteration) using step 1-7, and new outliers were identified.

*Identification of subtype-enriched genes*

Subtype-enriched gene analysis was carried out using the *scde* package with read counts table as input (Kharchenko et al., 2014). Error models for cells from different subtypes were fitted independently using the “groups” argument. For each subtype (except PSN-H), differential expression analysis was carried out with each of the other subtypes to generate 8 lists of upregulated genes (adjusted z score>1.7 and fold-change >2), and subtype-enriched genes were further identified by intersecting the lists. For PSN1-8, the lists of upregulated genes identified by comparison to PSN9 using *scde* were not used due to the low number of cells in PSN9; instead, the average log_2_(TMP+1) was used, and genes expressed ≥ 2-fold higher (Δlog_2_(TMP+1)>1) in PSN1-8 than PSN9 were included in the list. For PSN9, the enriched genes were identified by intersecting lists of upregulated genes generated by *scde* from all comparisons.

*Mouse lung scRNA-seq data analysis*

scRNA-seq data of the mouse lung was a combined dataset of two previous reports (Tabula Muris, 2018; Travaglini et al., 2020), containing 2153 cells from adult mice (postnatal 60-90 days) profiled using SMART-seq2 method. The previously assigned cell type annotations were used.

**Histology**

*Immunostaining on vagal ganglion sections*

Vagal ganglia were dissected freshly and fixed with 4% PFA at 4°C for 2 hours. After washing with PBS, ganglia were cryoprotected with 30% sucrose in PBS and embedded in optimal cutting temperature compound (OCT). Cryosections were cut at 18 µm and dried overnight at room temperature. After washing 2 x 5 min with PBS and 1 x 5 min with PBS+0.1% Triton X-100, sections were incubated with blocking solution (PBS+0.1% Triton X-100+5% normal donkey serum) for 30 min at room temperature, followed by primary antibodies at 4°C overnight. The next day, sections were washed 3 x 10 min with PBS+0.1% Triton X-100, incubated with secondary antibodies (1:500) for 45-60 min at room temperature. After washing again 3 x 10 min with PBS+0.1% Triton X-100 and then briefly rinsing in PBS, sections were mounted with fluoromount-G. Primary antibodies used are listed in the Key Resources Table.

*Single molecule fluorescent in situ hybridization (smFISH) with immunostaining*

Vagal ganglia from mice retrogradely labeled with WGA were mounted in OCT and freshly frozen on dry ice. Cryosections were cut at 12 - 14 µm and post-fixed for 45-60 min to preserve the WGA throughout *in situ* hybridization. smFISH (RNAscope) was performed using the supplier’s protocol (Advanced Cell Diagnostics, ACD), followed by the immunostaining protocol described above.

*Immunostaining on lung vibratome sections*

Mouse lungs were perfused with PBS through the pulmonary circulation and inflated with 2% low-melting point agarose in PBS. After removal from the body, the inflated lungs were fixed with 4% PFA in PBS for 5 hrs at 4°C. After washing thoroughly with PBS, lung lobes (left and right cranial) were separated and glued onto the cutting stage of the vibratome, and sections (350 µm) were cut parallel to the ventral surface of the lobes.

Vibratome sections were washed with PBS, blocked with PBS+0.5%Triton X-100+5% normal donkey serum for 1 hour at room temperature, then incubated with primary antibodies for 2.5 days at 4°C on a nutator/shaker. Antibody-stained sections were washed with PBS+0.5%Triton X-100 for 2 x 30 min, 2 x 1 hour, 1 x 2 hours, then incubated with secondary antibodies (1:500) for 1-2 days at 4°C on a nutator/shaker. Sections were washed as above after secondary antibody incubation, then post-fixed with 4% PFA in PBS for 1 - 2 hrs at 4°C, dehydrated with 50%, 75%, and 100% methanol, then cleared in BABB (Benzyl Alcohol:Benzyle Benzoate 1:2). Images were acquired with a Zeiss 780 confocal scanning microscope under 25x oil objective.

*Alkaline Phosphatase staining*

Mouse lungs were perfused and inflated as described above. After removal from the body, the inflated lungs were fixed with 4% PFA in PBS for 2 hours at 4°C. Whole lungs were then washed with PBS, incubated in PBS at 65-68°C for 2 hours to inactive endogenous alkaline phosphatase, and stained with BCIP/NBT solution (diluted in 0.1M Tris-HCl pH 9.6, 0.1M NaCl, 50mM MgCl2, 0.1% Triton X-100) overnight at room temperature. The next day, lungs were washed with PBS, post-fixed with 4% PFA in PBS overnight at 4°C, dehydrated with 50%, 75%, and 100% methanol, then cleared in BABB.

**Vagal ganglion injection**

Vagal ganglion injections were performed as previously described with minor modifications (Chang et al., 2015). Briefly, mice were intubated and kept anesthetized with 1.5% isoflurane (2-3% for induction) using a low-flow anesthesia system (Somnosuite, Kent Scientific, Physiosuite modules incorporated). To avoid complications from breathing movements, mice were ventilated (RoVent module) at 90 breaths per minute (10-12 cmH_2_O target pressure). The vagal ganglion was exposed from the ventral side of the neck. Injections were carried out using custom glass pipettes (pulled by micropipette puller P-87, Sutter Instrument) connected to UltraMicroPump (World Precision Instruments) with Micro4 controller (World Precision Instruments). Virus solution (200 nl), containing 0.05% Fast Green (Sigma, F7252) for visualization during injection, was injected to each ganglion at a rate of 20 nl per minute. For histological analysis, mice injected with AAV-CAG-DIO-tdTomato or AAV-Syn-DIO-hM3Dq-mCherry were sacrificed at least 3 weeks after ganglion injection, and mice injected with AAV-CMV-DIO-PLAP were sacrificed at least 2 weeks later.

**Chemical delivery**

*Trimethoprim*

Trimethoprim (TMP, Sigma T7883) was delivered to mice by oral gavage. On each day of treatment, a 50 mg/ml stock solution was freshly prepared by dissolving TMP in DMSO followed by vigorous vortexing. The stock solution was further diluted 1:5 in sterile water and immediately delivered in the appropriate volume at a dose of 300 µg/g body weight.

*Diphtheria toxin*

Diphtheria toxin (Biological Laboratory, Cat#150) was dissolved in PBS to make a 20 µg/ml stock solution and stored at -20°C. On each day of treatment, stock solution was diluted in sterile saline to make a 200 ng/ml injection solution and injected i.p. at a dose of 10 ng/g body weight.

**Breathing assays**

*Whole-body plethysmography*

Whole-body plethysmography recordings were performed on unanesthetized and unrestrained mice using Buxco chambers (450 ml, Model PY4211) or Vivoflow chambers (SciReq) and data was acquired and analyzed using usbAMP and iox2 software (SciReq). For ablation experiments, each mouse was pre-acclimated to the chamber for two consecutive days (minimum 1.5 hours per day) before the first day of recording (prior to DT injection) and for one day before the post DT recording (third day post DT injection). On each recording day, mice were recorded for at least 2 hours.

To analyze respiratory parameters, breaths in the first 30 min of recordings were not used, and breaths that had a tidal volume (TV) less than 0.05 ml or greater than 2 ml, or a peak inspiratory flow (PIF) less than 1 ml/s or greater than 25 ml/s were filtered out. To automatically select calm breaths, all breaths after filtering were segmented into 1 min windows, and the average change in PIF of every pair of consecutive breaths was calculated within all windows. The distribution of average PIF changes was calculated by the *density* function in R, and all 1-minute windows in the first peak (≤2x the first mode) were selected as quiet breathing periods (Figure S7E, F). Breaths in quiet breathing periods were used to plot distributions of individual parameters and calculate distribution modes or averages for comparison. Minute ventilation was calculated by summing up the TV of all calm breaths then dividing by the total time (in minutes).

*Inflation assay*

Mice were anesthetized by two consecutive i.p. injections of 20% urethane in saline (each 1 mg/g body weight) one hour apart, and surgery started one hour after the second injection. Additional injections(s) of 2-4 mg were given (total < 10mg) as needed if the toe pinch reflex was not completely abolished. Mice were intubated with a 1-inch 20G disposable catheter and connected to Physiosuite (Kent Scientific) supplied with room air at 50-60 ml/min. Body temperature was kept at 36°C using a heating pad (RightTemp, Kent Scientific). The trachea was exposed, and a suture was used to seal the endotracheal tube airtight in the trachea. Chest and abdominal skin were incised to expose the sternum and the left side of the diaphragm. Prior to opening the chest cavity, the ventilator was started in pressure control mode (RoVent, Kent Scientific, 150 breaths/min, target pressure: 10 cmH_2_O, PEEP: 2 cmH_2_O, I/E ratio: 1:1.5). The chest cavity was opened by cutting the sternum (along the midline to avoid bleeding). A two-lead stainless steel needle electrode was inserted into the left costal diaphragm, and diaphragm EMG signal was recorded with BioAmp connected to Powerlab (ADInstrument). Airflow in the ventilation circuit was monitored using a spirometer with a pneumotach flow head, and signals were recorded using Powerlab. Any mice that had diaphragm rhythmic activity when ventilating at 10 cmH_2_O, indicating an abnormal mechanoreflex, were not proceeded to inspiratory/expiratory challenge. Target pressure was then gradually lowered until diaphragm rhythmic activity initiated (6~8cmH_2_O) with a frequency equal to or higher than ventilation. Inflation and deflation holding were delivered by Physiosuite-RoVent for 5 s each. Between two consecutive holding challenges, there was at least 2 min of regular ventilation. Serial challenges on the same mouse followed an order that alternated between high pressures (>12 cmH_2_O) and low pressures (<12 cmH_2_O) to avoid atelectasis throughout multiple challenge trials.

Diaphragm EMG signals were high-pass filtered at 150 Hz. For each inspiratory challenge, the duration of individual EMG bursts (Ti) and the inter-burst interval (Te) were calculated. Changes in burst duration (ΔTi) and inter-burst interval (ΔTe) were determined by comparing values obtained during inspiration holding with those measured during expiration holding in the same animal.

**In vivo calcium imaging and activity analysis**

In vivo calcium imaging of the mouse vagal ganglion was performed as previously described ^20^. Briefly, retrograde labeling with WGA (WGA-Alexa Fluor 594 or WGA-CF594) was conducted 4-7 days prior to the imaging experiment. Mice were anesthetized with urethane following the same procedure used for the inflation assay and subsequently intubated with a 20G disposable catheter. The left vagal ganglion and nerve were exposed and carefully isolated from the carotid artery. The pharyngeal and superior laryngeal branches of the vagus nerve were transected, and the ganglion was separated from its central projections by a transection performed as superior as possible. The ganglion was then gently mobilized away from the jawbone and surrounding muscles and secured to a customized bar using Kwik-Sil silicone adhesive (WPI). To create a bath chamber around the ganglion, the neck skin was attached to a customized metal ring stabilized with a side bar. The chamber was filled with HEPES-buffered Krebs–Ringer solution, which also served as the immersion medium.

Imaging was performed with a Bruker Ultima 2Pplus multiphoton microscope equipped with a Chameleon Discovery dual-output laser and a 16× Nikon water-immersion objective. GCaMP fluorescence was excited at 920 nm, while WGA fluorescence was captured using either 810 nm (Alexa Fluor 594) or 920 nm (CF594) excitation. Images were acquired at 4 frames per second. Cells of interest were identified by co-localization of GCaMP and WGA signals. Regions of interest were manually extracted from the GCaMP images. Baseline fluorescence (F0) was defined as the mean signal intensity across 20 frames acquired prior to the onset of inflation/deflation challenges. Neural activity was quantified as ΔF/F0.

**Statistical analysis**

Sample sizes including number of ganglia and number of mice are reported for each experiment in figures or figure legends. Statistical analysis and plotting were performed using Graphpad Prism 10. All bar plots show the mean values ± SEM (error bars). Statistical methods for comparison are reported in figure legends. In figures, asterisks are used to denote significance, with * p < 0.05 and ** p < 10^-2^.
